# The *Parastagonospora nodorum* necrotrophic effector SnTox5 targets the wheat gene *Snn5* and facilitates entry into the leaf mesophyll

**DOI:** 10.1101/2021.02.26.433117

**Authors:** Gayan K. Kariyawasam, Jonathan K. Richards, Nathan A. Wyatt, Katherine Running, Steven S. Xu, Zhaohui Liu, Pawel Borowicz, Justin D. Faris, Timothy L. Friesen

## Abstract

*Parastagonospora nodorum*, causal agent of septoria nodorum blotch, is a destructive necrotrophic fungal pathogen of wheat. *P. nodorum* is known to secrete several necrotrophic effectors that target wheat susceptibility genes that trigger classical biotrophic resistance responses but resulting in susceptibility rather than resistance. SnTox5 targets the wheat susceptibility gene *Snn5* to induce necrosis. In this study, we used full genome sequences of 197 *P. nodorum* isolates collected from the US and their disease phenotyping on the *Snn5* differential line LP29, to perform genome wide association study analysis to localize the *SnTox5* gene to chromosome 8 of *P. nodorum*. *SnTox5* was validated using gene transformation and CRISPR-Cas9 based gene disruption. *SnTox5* encoded a small secreted protein with a 22 and 45 amino acid secretion signal and a pro sequence, respectively. The *SnTox5* gene is under purifying selection in the Upper Midwest but under strong diversifying selection in the South/East regions of the US. Comparison of wild type and SnTox5-disrupted strains on wheat lines with and without the susceptibility target *Snn5* showed that SnTox5 has two functions, 1) facilitating colonization of the mesophyll layer, and 2) targeting Snn5 to induce programmed cell death to provide cellular nutrient to complete its necrotrophic life cycle.

## Introduction

Plant pathogenic fungi secrete a variety of effectors that contribute to virulence during host infection. These effectors are secreted into the apoplast or internalized into the cytoplasm where they manipulate the host cell’s biological processes to promote host colonization (Lo Presti et al. 2015; Franceschetti et al. 2017). However, plants have evolved plant innate immunity including resistance (R) receptors that recognize effectors, resulting in effector triggered immunity (ETI) and pattern recognition receptors (PRRs) that recognize pathogen associated molecular patterns (PAMPs), resulting in PAMP-triggered immunity (PTI) (Jones and Dangle, 2006). Typically the resistance response results in programmed cell death (PCD) surrounding the infection site via an intense but localized reaction called a hypersensitive response (HR) that occurs along with, or as a result of a combination of physiological processes including, accumulation of reactive oxygen species (ROS), lipid peroxidation, ion fluxes, and deposition of callose on infection sites (Dodds and Rathjen, 2010; Balint-Kurti, 2019).

Localized PCD is highly effective against biotrophic fungal pathogens because this class of pathogens typically requires a living cell to extract nutrients. The PCD response is less effective against necrotrophic fungal pathogens because they thrive and acquire nutrients made available by cell death. Necrotrophic fungal pathogens release necrotrophic effectors (NEs), a group of effectors that target host genes to trigger necrosis, providing nutrients to the pathogen (Friesen and Faris, 2010; Faris and Friesen 2020). Unlike classical gene-for-gene interactions described for biotrophic pathosystems (Flor, 1971), necrotrophic interactions result in necrotrophic effector triggered susceptibility (NETS) (Liu et al. 2009). In the past two decades several NE-host susceptibility gene interactions have been described, including those found in the *Pyrenophora tritici-repentis*-wheat (Faris et al. 2013), *Pyrenophora teres*-barley (Liu et al. 2015), *Bipolaris sorokiniana*-wheat (McDonald et al. 2018), *C. victoriae*-oat, *C. victoriae*-Arabidopsis (Lorang et al. 2007), *Periconia circinata*-sorghum (Nagy and Bennetzen, 2008) and *Parastagonospora nodorum*-wheat pathosystems (Faris and Friesen 2020).

Four susceptibility genes targeted by necrotrophic effectors have been cloned. These include *Tsn1* (Faris et al. 2010), a wheat gene that confers sensitivity to SnToxA (Friesen et al. 2006), Ptr ToxA (Ciuffetti et al. 1997), and BsToxA (McDonald et al. 2018), the sorghum gene *Pc* that confers sensitivity to Pc-toxin (Nagy and Bennetzen, 2008), the *Arabidopsis* gene *Lov1* that confers sensitivity to victorin (Lorang et al. 2007), the wheat genes *Snn1* that confers sensitivity to SnTox1 (Shi et al. 2016), and *Snn3-D1* that confers sensitivity to SnTox3 (Zhang et al. 2021). Although Snn3-D1 has not been functionally shown to be involved in a resistance-like response, each of the other characterized susceptibility genes resemble genes involved in resistance responses to biotrophic pathogens, where *Tsnl*, *Pc*, and *Lov1* encode proteins with nucleotide binding (NB) and leucine rich repeat domains (LRR), and *Snn1* encodes a wall associated kinase (WAK) (Shi et al. 2016). These examples show that necrotrophic pathogens often hijack the plant’s resistance response system by targeting the defense response pathways to induce PCD, facilitating the completion of its pathogenic life cycle.

*P. nodorum* is a destructive necrotrophic fungal pathogen of wheat that causes the economically important disease septoria nodorum blotch (SNB). Extensive research over the last two decades has shown that *P. nodorum* deploys several proteinaceous necrotrophic effectors during the infection process. To date, nine such interactions have been identified including, SnToxA-*Tsn1* (Friesen et al. 2006), SnTox1-*Snn1* (Liu et al. 2004), SnTox267-*Snn2* (Friesen et al. 2007; Richards et al. 2021), SnTox267-*Snn6* (Gao et al. 2015), SnTox3-*Snn3-B1* (Friesen et al. 2008), SnTox3-*Snn3-D1* (Zhang et al. 2011; Zhang et al. 2021), SnTox4-*Snn4* (Abeysekara et al. 2009), SnTox5-*Snn5* (Friesen et al. 2012), and SnTox267-*Snn7* (Shi et al. 2015). Therefore, currently the wheat-*P. nodorum* system is recognized as a model for the study of the infection process of necrotrophic specialist pathogens (Oliver et al. 2012; Faris and Friesen 2020).

Currently three necrotrophic effector genes from the *P. nodorum* system have been cloned and functionally characterized, including *SnToxA*, *SnTox1*, and *SnTox3*. *SnToxA* is nearly identical to the *ToxA* gene found in *P. tritici-repentis* and *Bipolaris sorokiniana* and it encodes a 13.2 kDa mature protein that targets *Tsn1* indirectly to cause necrosis (Ciuffetti et al. 1997; Friesen et al. 2006; McDonald et al. 2018). *SnTox3* was the second *P. nodorum* necrotrophic effector gene cloned (Liu et al. 2009), encoding for a mature 17.5 kDa protein that targets *Snn3-B1* (Friesen et al. 2008) and *Snn3-D1* (Zhang et al. 2011; Zhang et al. 2021) to induce necrosis. SnTox3 also functions to suppress the host defense through an direct interaction with TaPR1 proteins (Breen et al. 2016; Sung et al. 2021). The most recent necrotrophic effector gene that was cloned in this system was *SnTox1* that encoded a 10.3 kDa protein that targets Snn1 directly (Shi et al. 2016) to trigger an oxidative burst, upregulation of PR-genes, and DNA laddering (Liu et al. 2012). Liu et al. (2016), showed that in addition to targeting Snn1, SnTox1 had the ability to bind chitin and protect the fungal cell wall from wheat chitinases.

SnTox5 is also a proteinaceous necrotrophic effector and interacts with the susceptibility gene *Snn5* (Friesen et al. 2012). *Snn5* was mapped to chromosome 4B in the Lebsock×PI94749 (LP749) population using culture filtrates of *P. nodorum* isolate Sn2000 that contained the SnTox5 protein (Friesen et al. 2012). Susceptibility to Sn2000 also mapped to the *Snn5* locus on 4B showing that *P. nodorum* was using the necrotrophic effector SnTox5 as a virulence factor to target the *Snn5* susceptibility gene to induce PCD resulting in disease.

Here we used whole genome sequencing of 197 *P. nodorum* isolates in a genome wide association study (GWAS) to identify, clone, and functionally validate the *SnTox5* gene from *P. nodorum* isolate Sn2000. We also used laser confocal microscopy to characterize the role that SnTox5 plays in *P. nodorum* leaf colonization. This study provides further characterization of how *P. nodorum* is using its arsenal of necrotrophic effectors to target the host defense pathways to complete its pathogenic life cycle.

## Materials and methods

### Disease phenotyping

A set of 197 *P. nodorum* isolates was collected from geographically diverse winter, spring, and durum wheat growing regions of the US. The population consisted of 51 isolates collected from spring wheat in North Dakota and Minnesota, 45 isolates collected from durum wheat in North Dakota, nine isolates collected from winter wheat in South Dakota, and 92 isolates from winter wheat regions of the United States representing Arkansas, Georgia, Maryland, New York, North Carolina, Ohio, Oklahoma, Oregon, South Carolina, Tennessee, Texas, and Virginia, (Richards et al. 2019). Culture preparation and phenotyping was done as described by Friesen and Faris, (2012). In brief, a dried agar plug of each isolate was place on V8-PDA (150 ml of V8 juice, 10 g of Difco potato dextrose agar, 3g of CaCO_3_, 10g of Agar in 1000 ml of water) and allowed to rehydrate for 15 minutes. The rehydrated plug was then streaked across the plate to evenly distribute the spores and the plate was incubated at room temperature under continuous light for seven days or until pycnidia emerged. Plates with pycnidia were flooded with sterile-distilled water and agitated with a sterile inoculation loop to stimulate the release of pycnidiospores. Spores were harvested, and the spore concentration was adjusted to 1 × 10^6^ spores/mL and two drops of Tween20 were added per 100 ml of spore suspension.

All isolates were phenotyped for disease reaction on LP29, the differential line for *Snn5*, which is the sensitivity gene targeted by SnTox5 (Friesen et al 2012). LP29 is a progeny line chosen from the doubled haploid population derived from the cross Lebsock × PI94749 that segregated for the wheat susceptibility genes *Snn5* and *Tsn1* (Friesen et al 2012). Each replicate consisted of a single cone with three plants of LP29. Borders were planted with the wheat cultivar Alsen to reduce any edge effect. Plants were grown for approximately fourteen days. Plants at the two-to-three leaf stage were inoculated with a spore suspension using a pressurized paint sprayer. Leaves were inoculated until runoff and kept in a lighted mist chamber at 100% relative humidity at ∼21 °C for 24 hours prior to being moved into a climate-controlled growth chamber at 21 °C with a 12-hour photoperiod for six additional days. At 7 days post-inoculation, disease was evaluated using a 0-5 rating scale based on the lesion type as described in Liu et al. (2004). Each experiment was performed in three replications and the average of the three replicates was used in downstream analysis.

### Whole genome sequencing and variant identification

Raw sequencing reads for each isolate were generated using the Illumina HiSeq 4000 platform at BGI Americas Corp and uploaded to the NCBI short read archive under BioProject PRJNA398070 (Richards et al. 2019). Raw sequencing reads were trimmed using Trimmomatic v0.36 (Bolger et al. 2014) and were mapped to the reference genome sequence of *P. nodorum* isolate Sn2000 using BWA-MEM (Li, 2013). SAMtools ‘mpileup’ (Li et al.2009) was used to identify SNPs/InDels and the variants were filtered based on the genotype quality where only the polymorphisms with genotype quality equal to or greater than 40 with the support of a minimum of three reads were used for downstream analysis. All heterozygote calls were marked as missing data and variants with 30% or more missing data were removed from the dataset. In addition, markers with a minor allele frequency of less than 5% were filtered out from the final dataset used for genome-wide association study analysis.

### Genome-wide association study (GWAS) analysis

Mapping for GWAS was performed using GAPIT (Lipka et al. 2012; Tang et al. 2016) and TASSEL v5 (Bradbury et al. 2007). For the association mapping conducted using TASSEL v5, a naïve model and a model comprised of the first three components of PCA as fixed effects were used. For the analysis performed with GAPIT, models with a kinship matrix (K) using EMMA as a random effect and models using a combination of both PCA and K were used. The most robust model was selected based on Q-Q plot results. A Bonferroni correction was used to adjust the *P*-value in the R statistical environment and the markers were considered significant when an adjusted *P*-value was equal to or less than 0.05.

### Identification of candidate genes

The candidate region for *SnTox5* in the Sn2000 genome was identified using GWAS analysis. The region was screened for genes encoding small secreted proteins using SignalP v4.1 (Petersen et al. 2011) and EffectorP v1.0 (Sperschneider et al. 2016). A gene encoding a small secreted protein that contained the marker with the most significant marker-trait association from GWAS analysis was considered the top candidate for *SnTox5*.

### Deletion of SnTox5 in the virulent isolate Sn2000

The disruption of *SnTox5* was carried out using a CRISPR-Cas9 ribonucleoprotein-mediated technique as described in Foster et al. (2018). In brief, FASTA sequence of *SnTox5* in Sn2000 was input into E-CRISP at http://www.e-crisp.org/E-CRISP/ to select the primer template for the sgRNA. Oligonucleotides were purchased from Eurofins Genomics, KY. The sgRNA was synthesized using the sgRNA synthesis kit NEB#E3322 from New England Biolabs following the manufacturer’s protocol. The resulting sgRNA was purified prior to complexing with Cas9-NLS via the RNA clean and concentrator -25 kit from Zymo Research following the manufacturer’s instructions.

Primers, Tox5HygDonor F1 and Tox5HygDonor R1 (Supplementary Table 1) were designed to amplify the complete hygromycin resistance gene as the donor DNA. Each primer consisted of a 40 bp sequence homologous to the flanking region adjacent to the protospacer adjacent motif (PAM) site and 3 bp upstream of the PAM site that is incorporated into the ends of the hygromycin resistance gene, *cpc-1:hyg^R^*, which was amplified from the pDAN vector (Liu et al. 2012) as the template.

Fungal protoplast generation and transformation were performed as described in Liu and Friesen (2012). Cas9-NLS were complexed with sgRNAs and then mixed with the donor DNA that was transformed into protoplasts of Sn2000. Protoplasts were plated on regeneration medium agar supplemented with hygromycin B. Regenerated colonies were picked and screened for *SnTox5* disruption using the primers SnTox5_pENTR_F1_bac and SnTox5_pENTR_R1 (Supplementary Table 1) and for the presence of the hygromycin resistance gene. Two *SnTox5-*disrupted strains and one strain with an ectopic insertion of the hygromycin resistance gene were used for downstream phenotypic analysis on the host.

### QTL analysis of the LP749 population using SnTox5 gene disruption strains

The LP749 population (Friesen et al 2012) was used to map the SnTox5-*Snn5* interaction. The same population was inoculated separately with the two *SnTox5* gene disruption strains Sn2kΔTox5-10 and Sn2kΔTox5-15, the ectopic strain Sn2k-ect7, and the wild type strain Sn2000. Side by side inoculations using each of the four strains were completed on full LP749 populations and the disease was evaluated at seven days post-inoculation as described above.

Averages disease reactions from three replicates were used to perform composite interval mapping (CIM) to evaluate the significance of the SnTox5-*Snn5* and SnToxA-*Tsn1* interactions for the inoculation of each *P. nodorum* strain using Qgene v4.4.0 (Joehanes and Nelson, 2008). A permutation test that consisted of 1000 iterations yielded a LOD threshold of 3.0 at an experimental-wise significance level of 0.05 and was used to evaluate the significance of the resulting QTL.

### Expression of SnTox5 in the avirulent P. nodorum isolate Sn79-1087

The Gateway cloning system (Gong et al. 2015) was used to develop constructs with *SnTox5*. Approximately 1.7 kb of the genomic region of *SnTox5*, including a 1 kb region upstream of the gene that included the putative promotor region (Supplementary Figure 1) was amplified with forward primer SnTox5_DONOR_F and reverse primer SnTox5_DONOR_R (Supplementary Table 1). Each primer consisted of a full length attB sequence at the 5’ end. The PCR amplicon with an attB sequence at the end was visualized using gel electrophoresis and purified using the GeneJet Gel Extraction Kit (Thermo Scientific). Fragments were cloned into the pDONOR vector via a BP Clonase reaction (Invitrogen). The pDONOR vector, containing the resistance gene zeocin, was transformed into *E. coli* and transformed colonies were selected on low salt Luria-Bertani broth (LB) agar medium (10g of tryptone, 5g of NaCl, 5g of yeast extract, and 16g of agar in 1000 ml of water) amended with zeocin (50 µg/ml). Five *E. coli* transformants were picked and inoculated in 2 ml of low salt LB with zeocin and used to extract plasmid using the Monarch plasmid miniprep kit (New England Bio Labs). Presence of the genomic fragment containing *SnTox5* was verified by Sanger sequencing using the Tox5_Seq_F, M13 forward and reverse primers (Supplementary Table 1). The extracted pDONOR plasmid with the insertion was used to perform an LR Clonase reaction as instructed by the manufacturer (Invitrogen) to transfer the genomic region into the destination vector, pFPL-RH, that contained the hygromycin resistance cassette. The construct was linearized using *Pme*I and concentrated to 1 µg/µl. Fungal protoplasting and transformation was performed as described in Liu and Friesen (2012) to transform Sn79-1087 (avirulent isolate) with *SnTox5*. Colonies that developed on regeneration media with hygromycin (100 µg/ml) were screened for the presence of the gene through PCR amplification using primers SnTox5_PENTR_F and SnTox5_PENTR_R (Supplementary Table 1). Two transformants, Sn79+Tox5-3 and Sn79+Tox5-4 and wildtype Sn79-1087 were inoculated onto the LP749 population and QTL analysis was done as mentioned previously. Furthermore, culture filtrates of Sn79+Tox5-3 and Sn79-1087 were prepared as described (Liu et al. 2004) and used to infiltrate the LP749 population. Sensitivity was scored using a 0-3 rating scale where 0 was rated as insensitive and 3 was rated as highly sensitive (Friesen and Faris 2012). Data was used for QTL analysis as described previously.

### Homology between SnTox3 and SnTox5

Protein BLAST was performed against the NCBI non-redundant protein sequence database using the amino acid sequence of SnTox5 as the query. SnTox3, which was the protein with highest homology to SnTox5, was aligned to SnTox5 using “Geneious Alignment” option in Geneious Prime. In addition, disulfide bridges in SnTox5 that formed between cysteine residues were predicted using the web-based application DiANNA1.1 (http://clavius.bc.edu/~clotelab/DiANNA/).

### Population genetics and haplotype analysis of SnTox5

BAM files for 197 isolates of the GWAS panel were developed as described above and were used to extract reads mapped to chromosome 8:53219-53872bp of the Sn2000 genome using SAMtools (Li et al. 2009) and a BED file, creating FastQ files for each isolate. *De-novo* assembly of *SnTox5* for each isolate was completed using SPADES v.3.11.1 (Nurk et al. 2013) with default settings and *SnTox5* sequences for each isolate were developed for use in haplotype analysis. In addition, coverage of the *SnTox5* gene in each isolate was calculated using the ‘coverage’ function of BEDTools (Quinlan et al. 2010). Isolates with more than 50% of the *SnTox5* gene were considered to have the gene, whereas, isolates with coverage less than or equal to 50% were considered to lack the gene. Genomic sequences of *SnTox5* for isolates that contained complete coverage of the gene were converted to FASTA format and imported into DNASP v6 for population genetic analysis. The predicted SnTox5 amino acid sequences for each haplotype were aligned using the web-based sequence aligner MULTalign (Corpet, 1988) to identify amino acid sequence variation of the isoforms of SnTox5.

### Statistical analysis of variation in disease caused by different isoforms of SnTox5

Analysis of variance (ANOVA) and Fisher’s least-significant difference (LSD) were calculated to compare virulence of SnTox5 isoforms that were harbored by more than ten isolates based on the average disease reaction on LP29. In addition, isolates harboring fourteen active isoforms were grouped based on the amino acid residues at the 155^th^ and 156^th^ position of SnTox5, and ANOVA and LSDs were calculated for each group based on the average disease reaction on LP29. To account for the unbalanced sample size, a type III ANOVA was calculated using the package ‘car v3.0-10’ (Fox and Weisberg, 2019) and the resulting sums of square was used to calculate the LSDs at 0.05 experimental significance level using the package ‘agricolae v1.33’ (De Mendiburu, 2009) in the R programming environment in both analyses.

### Temporal expression profile of SnTox5 during the infection process

Secondary leaves of 14-day old ‘Lebsock’ were inoculated with Sn2000 and three samples of leaf tissue were collected at 4, 12, 24, 48, 72, 96, and 120 hours post-inoculation (hpi). Total RNA was extracted from leaves and purified using the RNeasy plant mini kit (Qiagen) according to the manufacturer’s protocol. RNA was quantified using a Qubit and 300 ng of total RNA from each time point was used to develop cDNA using the GoScript^TM^ Reverse Transcription System (Promega). With the use of gene specific primers SnTox5_qPCR_F and SnTox5_qPCR_R, qPCR was performed for all timepoints with three biological and three technical replicates. The *P. nodorum* actin gene was amplified as the internal control using previously published primers (Liu et al. 2009) (Supplementary Table 1).

### Laser confocal microscopic analysis of the infection process involving the SnTox5-Snn5 interaction

A construct with the *mCherry* gene coupled to the promoter of *SnTox1* was cloned into the pFPL-Cg vector containing a geneticin resistance cassette using the Gateway cloning system (Gong et al. 2015). The construct was linearized using *Pme*I and concentrated to 1 µg/µl. *P. nodorum* strains Sn2000, Sn2kΔTox5-15, Sn79+Tox5-3 and Sn79-1087 were transformed with the linearized construct, as explained in Liu and Friesen (2012), and transformants were inoculated onto two-week-old plants of the SnTox5 differential wheat line LP29. In addition, Lebsock was inoculated with *P. nodorum* strains Sn2000 and Sn2kΔTox5-15. Two leaf samples were collected at 4, 12, 24, 48, 72, 96, and 120 hpi and a 2.5 cm-long cutting from the middle portion of each secondary leaf was placed on a glass slide. Ecomount mounting media (Biocare Medical, CA) was applied to the sample and a coverslip was placed on the sample without introducing any air bubbles. The slides were dried overnight under room temperature prior to preservation at 2-8 °C. Three such replicates were conducted for each isolate-wheat line combination.

All the prepared slides were observed under an LSM700 laser scanning confocal microscope using 20x and 40x objectives where images captured under 20x were used for calculation of fungal mass and images captured under 40x were used to characterize the features of the infection process. Two different channels were used where the red channel ((Ex555/Em 639 nm) was assigned to capture the fluorescence emitted by mCherry and the green channel (Ex488/Em555 nm) was used for auto fluorescent detection of the leaf structure (Zeiss Thronwood, NY). To observe the infection process in different tissues within the leaf, Z-stack images were taken at different depths of the leaf from upper to lower epidermis with the use of ZEN.v. 11(Zeiss Thronwood, NY). The 2-D images were processed using Imaris v9.6 software (Bitplane, CT) for microscopic characterization and 3-D images were reconstructed using Imaris v9.6 for the volume analysis. Animated figures were created using the web-based application BioRender (BioRender.com). Average fungal volume was calculated after constructing the surface of the inoculated wheat leaf sample at a minimum of two infection sites per sample at each timepoint in one experimental replicate. Three such replicates were conducted, meaning the average volume was calculated for six infection sites per time point under the 20x objective lens.

### Development of Snn5 mutants of LP29 and microscopic analysis of infection caused by Sn2000 and Sn2kΔTox5-15

The *Snn5* differential line, LP29, which carries the *Snn5* allele from Lebsock, was used for mutagenesis to generate LP29ems lines. LP29 seeds were treated with 0.25% ethyl methanesulfonate (EMS) in 0.05 M phosphate buffer as described in Williams et al. (1992). The M_2_ generation was infiltrated with Sn2000K06-1 (Friesen et al. 2006) culture filtrates containing SnTox5. Ten to fourteen M_2_ individuals per M_1_ were evaluated. Plants were scored for presence or absence of necrosis 5 days after infiltration. SnTox5-insensitive mutants were self-pollinated to obtain M_3_. LP29 M_3_ families were infiltrated with Sn79+Tox5-3 culture filtrates to confirm insensitivity. M_3_ and M_4_ plants from the LP29 mutant line LP29ems931, hereafter designated LP29Δ*snn5* were used in this study.

## Results

### Genome-wide association study (GWAS) identifies a SnTox5 candidate gene

To identify candidate genes *SnTox5*, an association mapping approach was used to identify significant marker-trait associations in the *P. nodorum* genome using a natural population of 197 *P. nodorum* isolates. The genotypic data for the GWAS analysis was generated by aligning the whole genome sequences of 197 *P. nodorum* isolates to the SnTox5-producing Sn2000 reference genome (Richards et al. 2018). A total of 1,026,859 SNPs and insertion/deletions were identified and after filtering, a final set of 402,601 high confidence markers were used in GWAS analysis. The 197 *P. nodorum* isolates were phenotyped on LP29, where the average disease reaction caused by each isolate ranged from 0 to 4.33 (Supplementary Table 2). Both genotypic and phenotypic data were used to perform GWAS analysis using both GAPIT and TASSEL v5 applications and the most significant marker trait association was identified for a SNP (*P*=6.71E-11) on chromosome 8 at the 53,300bp position. The most significant SNP resided in the gene *Sn2000_06735* and therefore this gene became our *SnTox5* candidate (Figure 1).

**Figure 1.**
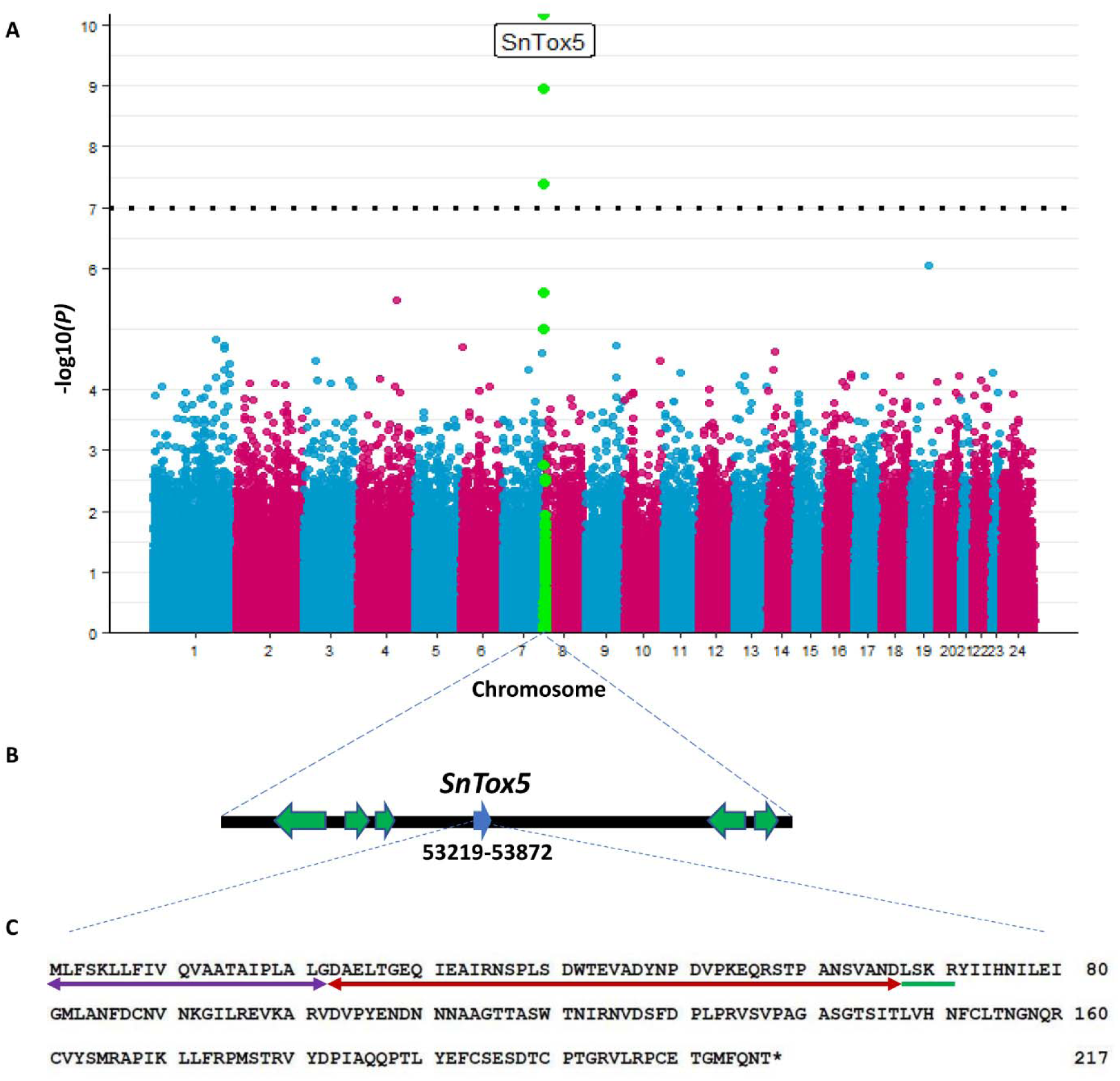
Genome-wide association mapping (GWAS) analysis of *SnTox5*. **A**) Manhattan plot of the GWAS performed using the phenotypic data from each isolates of the United States *Parastagonospora nodorum* population on differential line LP29. The x-axis labels denote chromosome numbers of the *P. nodorum* genome and the y-axis represents the -log10 transformation of the *P*-value for significance of the marker trait association. The horizontal dotted line represents the Bonferroni significance threshold at the 0.05 level of probability. **B)** The genomic location of *SnTox5* using the Sn2000 genome sequence as a reference. **C).** Amino acid sequence of SnTox5 from Sn2000. Purple and red double headed arrows represent the predicted signal peptide and the pro-sequence, respectively. The green line represents the putative Kex2 protease site and the “*” represents the stop codon.

The *Sn2000_06735* gene spanned from 53,219 to 53,872 bp on chromosome 8 of the Sn2000 genome (Figure 1) and was a 654bp intron-free gene with a putative TATA box 171 bps upstream of the start site (Supplementary Figure 1). The gene encoded a small secreted protein consisting of 217 amino acids with the first 22 amino acids predicted to be a signal peptide. A putative Kex2 protease site was identified at the 67^th^ amino acid, marking a putative 45 amino acid pro-sequence following the signal peptide. Sn2000_06735 also contained six cysteine residues predicted by DiANNA1.1 (http://clavius.bc.edu/~clotelab/DiANNA/) to form three di-sulfide bridges (Figure 1). In addition, BLASTp analysis against the NCBI database showed that SnTox5 had 45.13% homology to SnTox3, and pairwise alignment between the two protein sequences showed that the six cysteine residues were conserved, suggesting that Sn2000_06735 had both sequence and structural similarity to SnTox3 (Supplementary Figure 2).

### Deletion of Sn2000_06735 converts virulence to avirulence in the presence of Snn5

To validate that *Sn2000_06735* was *SnTox5*, *Sn2000_06735* was disrupted in *P. nodorum* isolate Sn2000 by inserting the hygromycin resistant cassette (*hyg^R^*) into *Sn2000_06735* using a CRISPR-Cas9 mediated gene disruption. The gene was successfully disrupted in five out of 24 transformants evaluated. Two disruption isolates designated Sn2kΔTox5-10 and Sn2kΔTox5-15 as well as an isolate with an ectopic insertion of *hyg^R^* designated Sn2k-ect7 were used for further analysis.

Sn2kΔTox5-10, Sn2kΔTox5-15, Sn2k-ect7 and the wild type isolate Sn2000 were inoculated onto LP29, the differential line for *Snn5*. Both Sn2000 and Sn2k-ect7 were able to induce typical necrotic lesions (Figure 2A). However, the two *Sn2000_06735* disrupted strains failed to cause necrosis on LP29 (Figure 2A). This suggested that Sn2000_06735 was targeting *Snn5* to cause disease, therefore *Sn2000_06735* will hereafter be referred to as *SnTox5*.

**Figure 2.**
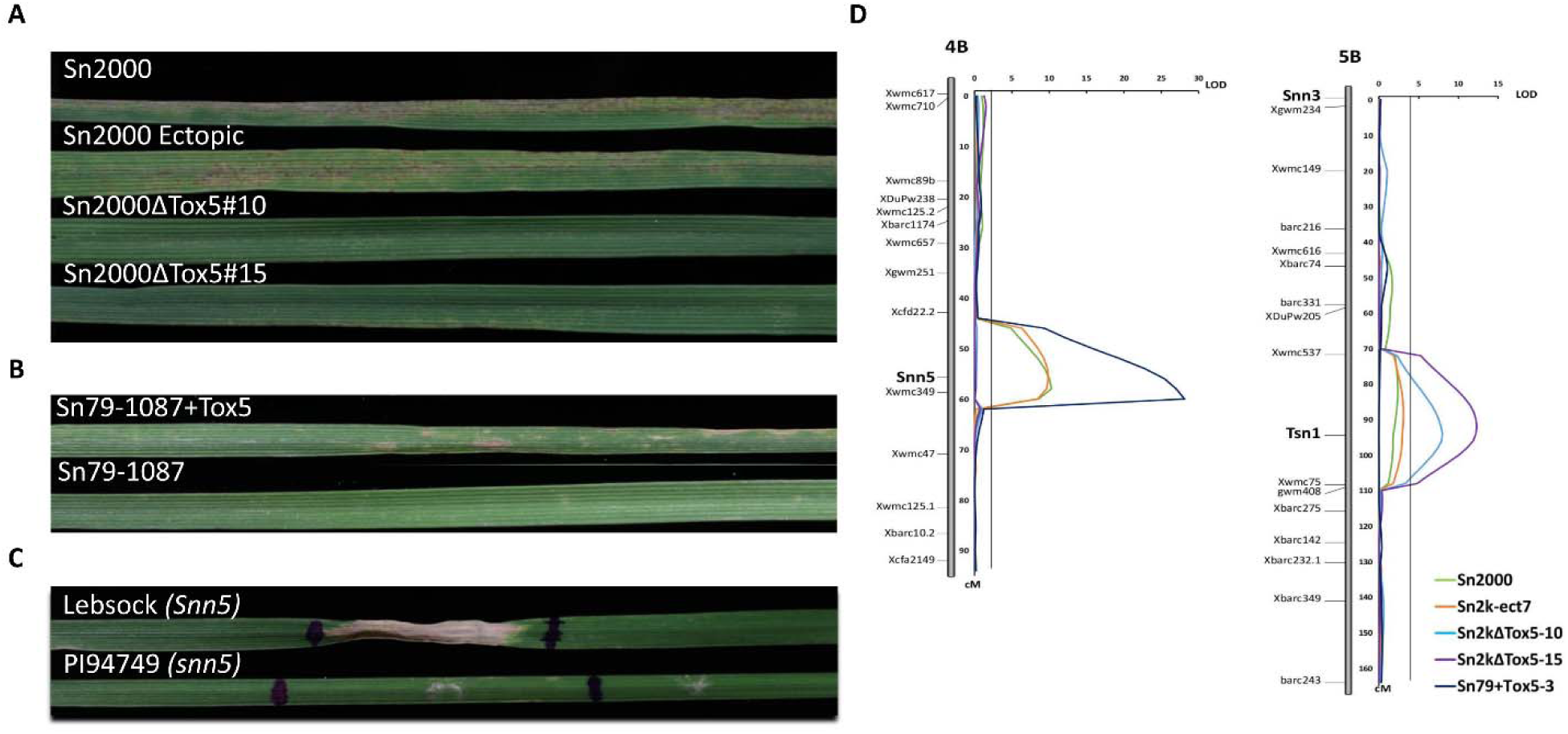
Phenotypic and QTL analysis validation of *SnTox5*. **A)** Phenotype of LP29 (*Snn5*) inoculated with Sn2000, Sn2k-ect7, and Sn2000 *SnTox5* gene-disruption mutants Sn2kΔTox5-10 and Sn2kΔTox5-15. **B)** Phenotype of LP29(*Snn5*) inoculated with the avirulent isolate Sn79-1087 (bottom) and the gain-of-function transformant Sn79+Tox5-3 (top). **C)**. Infiltration of culture filtrate of Sn79+Tox5-3 on the parental lines of the LP749 population including Lebsock (top) and PI94749 (bottom). **D)** QTL analysis on the LP749 population using strains Sn2000, Sn2k-ect7, Sn2kΔTox5-10, Sn2kΔTox5-15, and Sn79+Tox5-3, illustrating the significance of *Tsn1* and *Snn5* in the presence and absence of the *SnTox5-Snn5* interaction.

*Snn5*, the susceptibility target for SnTox5, was originally mapped using the LP749 double haploid population infiltrating culture filtrates containing SnTox5 (Friesen et al. 2012). The LP749 population was therefore inoculated with Sn2kΔTox5-10, Sn2kΔTox5-15, Sn2k-ect7, and the wild type isolate Sn2000. A significant QTL, previously described by Friesen et al. (2012), was identified on chromosomes 4B at the *Snn5* locus for both the wild type and ectopic strains (Figure 2D). In our experiment, the *Snn5* locus explained 33% and 32% of the variation in disease with LOD values of 10.31 and 9.65 for the disease caused by Sn2000 and Sn2k-ect7, respectively (Figure 2D, Table 1). In addition, the *Tsn1* locus explained 10% of the variation in disease caused by Sn2000 and 11% of the variation in disease caused by Sn2k-ect7 (Table 1). The significance of the *Snn5* locus was eliminated for the two *SnTox5* gene disruption mutants. As would be expected in the absence of the SnTox5-Snn5 interaction, the significance of *Tsn1* increased for both the mutants, explaining 26%-36% of the phenotypic variation with LOD values ranging from 7.82 to 12.3 (Figure 2D, Table 1).

**Table 1.** Composite interval mapping (CIM) analysis of QTL associated with Sn2000 (wild type), Sn2k-ect7 (isolate with an ectopic insertion of the hygromycin resistance gene, *cpc-1:hyg^R^*), Sn2kΔTox5-10, Sn2kΔTox5-15 (*SnTox5* disruption mutants of Sn2000) and Sn79+Tox5-3 (Sn79-1087 transformed with SnTox5) for the inoculation of the LP749 double haploid wheat population derived from the Lebsock (*Snn5*) × PI94749 (*snn5*) cross.

| Isolate | LOD<sup>□</sup> |  | R<sup>2</sup> |  |
| --- | --- | --- | --- | --- |
|  | *Snn5* | *Tsn1* | *Snn5* | *Tsn1* |
| Sn2000 | 10.31* | 2.37 | 0.33 | 0.10 |
| Sn2k-ect7 | 9.65* | 3.03* | 0.32 | 0.11 |
| Sn2kΔTox5-10 | 0.21 | 7.82* | 0.01 | 0.26 |
| Sn2kΔTox5-15 | 0.14 | 12.30* | 0.01 | 0.36 |
| Sn79+Tox5-3 | 27.00* | 0.15 | 0.66 | 0.01 |

<sup>□</sup> Permutation test with 1000 iterations resulted in a LOD threshold of 3.00 at *P*=0.05.
The “\*” represents significant QTL.

### Insertion of SnTox5 converts avirulence to virulence in the presence of Snn5

*P. nodorum* isolate Sn79-1087 is avirulent on LP29, and the *SnTox5* gene is completely absent (Figure 2B). Transformation of Sn79-1087 with *SnTox5*, along with its native promoter, converted the transformed strains Sn79+Tox5-3 and Sn79+Tox5-4 into virulent strains on LP29 (Figure 2B). Sn79+Tox5-3 caused an average disease reaction of 3.0 on Lebsock and an average disease reaction of 0 on PI94749, whereas Sn79-1087 (wild-type) showed no disease on either line.

The LP749 population segregated for disease caused by Sn79+Tox5-3 and subsequent analysis revealed a QTL at the *Snn5* locus with a LOD value of 27.00, explaining 66% of the disease variation (Figure 2D, Table 1). No other QTL were identified for this transformant, showing that SnTox5 was sufficient to cause disease in the presence of *Snn5*. The QTL analysis data from the LP749 population using the *SnTox5* disruption mutants and the *SnTox5* gain-of-function transformants further validated that *Sn2000_06735* was *SnTox5*.

### Isoform diversity of SnTox5 varies across geographical regions

The 197 *P. nodorum* isolates were collected from spring, winter, and durum wheat producing regions of the US and were screened for presence/absence of *SnTox5* using whole genome resequencing. Of the 197 isolates, 149 (75.6%) contained greater than 50% of the *SnTox5* gene and 48 (24.4%) isolates lacked the gene. However, the possibility of false negative results due to the low sequencing coverage of the region cannot be completely ignored. Of the 149 isolates that harbored the *SnTox5* gene, 128 had a complete gene sequence with sufficient sequence coverage and were therefore used for further haplotype analysis. The 128 isolates consisted of 22 nucleotide haplotypes resulting from polymorphisms at 118 nucleotide positions with an overall haplotype gene diversity (H_d_) of 0.861, indicating high levels of diversification of *SnTox5* within the US natural population. Haplotypes 17,18, 19, 20, 21, and 22, which were found in eleven of the isolates, contained a premature stop codon (Table 3). These haplotypes with a premature stop codon accounted for 105 polymorphic sites where the nature of polymorphisms indicated the presence of repeat induce polymorphism within *SnTox5*.Of the remaining 117 isolates with functional *SnTox5* haplotypes, 47 were found in the Upper Midwest population (44.76% of the Upper Midwest isolates), 47 were found in the South/East population (70.15 % of the South/East isolates), 7 were found in the Oregon population (87.5 % of Oregon isolates), and 17 were found in the Oklahoma population (100 % of Oklahoma isolates).

**Table 2.** Calculation of pN/pS ratios for the entire *SnTox5* gene and the region of the gene that encodes for the mature protein in the Upper Midwest and South/East populations of *P. nodorum*.

|  | Upper Midwest (n=47) |  | South/East (n=47) |  |
| --- | --- | --- | --- | --- |
|  | Entire gene | Mature protein encoding region<sup>a</sup> | Entire gene | Mature protein encoding region<sup>a</sup> |
| Synonymous SNPs | 2 | 2 | 1 | 1 |
| Synonymous Sites (average) | 154.55 | 108.22 | 156.97 | 107.63 |
| Nonsynonymous SNPs | 6 | 3 | 7 | 6 |
| Nonsynonymous Sites (average) | 487.45 | 341.78 | 494.03 | 342.37 |
| pN/pS (sites/average sites)<sup>b</sup> | 0.95 | 0.47 | 2.22 | 1.89 |

<sup>a</sup>base pairs 201 to 654 of the *SnTox5* encodes for the mature protein. The sequence resulted from the cleavage of signal peptide and pro-domain was considered for the calculation of pN/pS ratio.
<sup>b</sup> pN/pS ratios were calculated using the equation , $pN/pS = (\text{nonsynonymous SNPs} /$ $\text{nonsynonymous sites (average)}) / (\text{synonymous SNP} / \text{synonymous sites (average)})$ . $pN/pS < 1$ indicates that the *SnTox5* is undergoing purifying selection and $pN/pS > 1$ indicates that the gene is undergoing diversifying selection in the population.

**Table 3.**
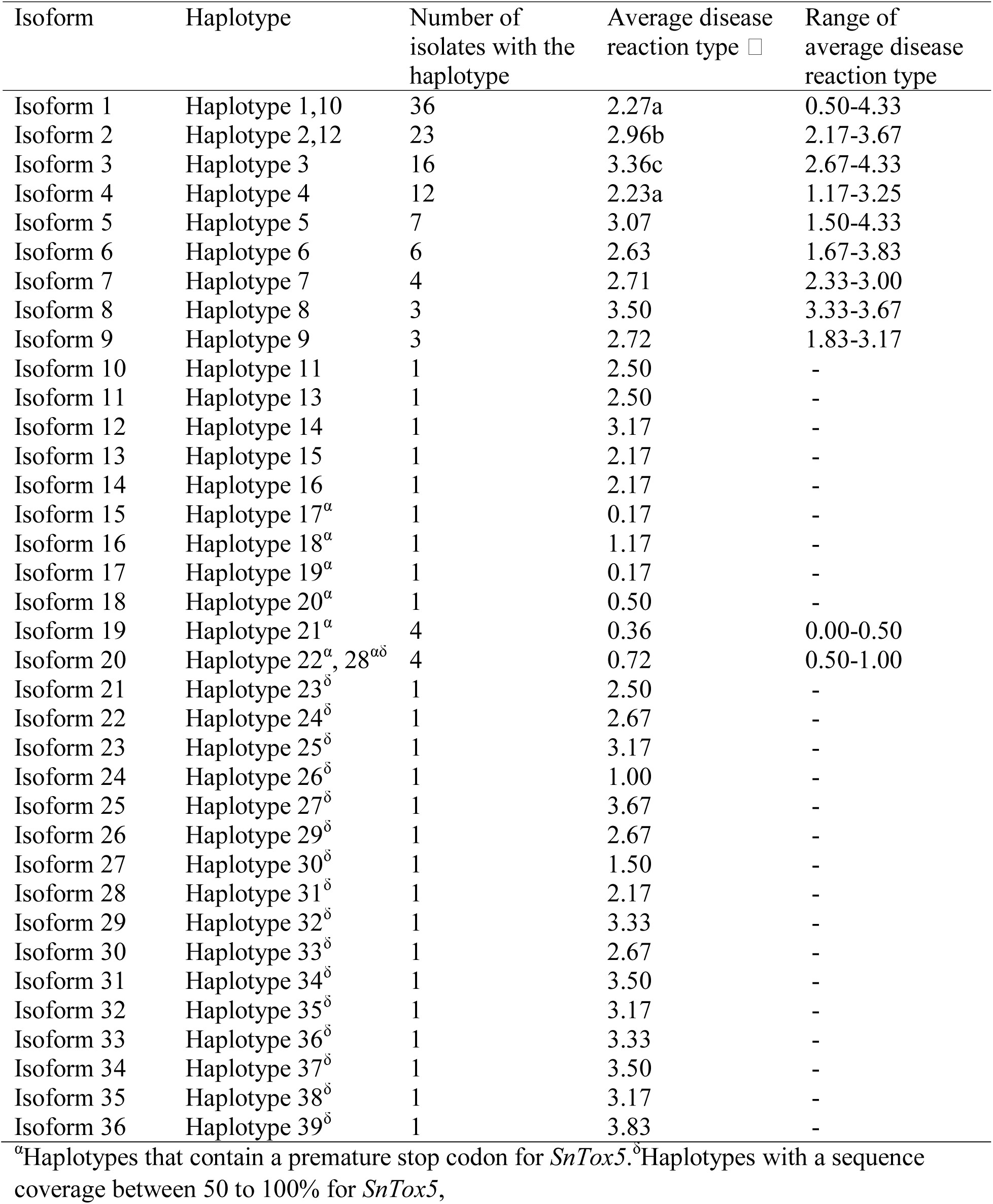
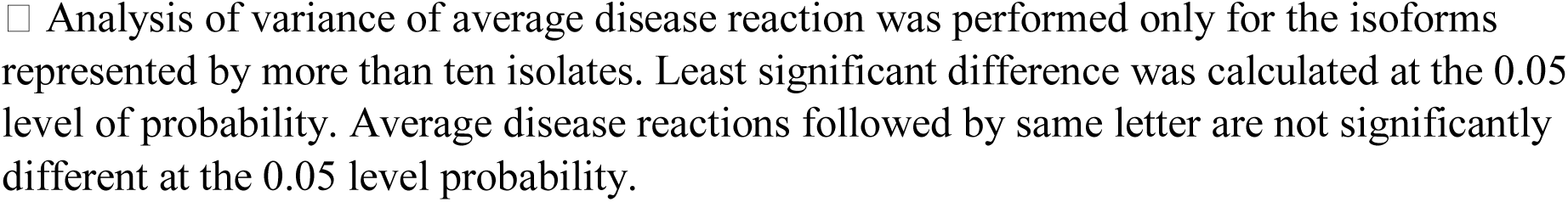
Average disease reaction type of isolates producing the different isoforms of SnTox5 on LP29 with in the United States population of *P. nodorum*.

| Isoform | Haplotype | Number of isolates with the haplotype | Average disease reaction type □ | Range of average disease reaction type |
| --- | --- | --- | --- | --- |
| Isoform 1 | Haplotype 1,10 | 36 | 2.27a | 0.50-4.33 |
| Isoform 2 | Haplotype 2,12 | 23 | 2.96b | 2.17-3.67 |
| Isoform 3 | Haplotype 3 | 16 | 3.36c | 2.67-4.33 |
| Isoform 4 | Haplotype 4 | 12 | 2.23a | 1.17-3.25 |
| Isoform 5 | Haplotype 5 | 7 | 3.07 | 1.50-4.33 |
| Isoform 6 | Haplotype 6 | 6 | 2.63 | 1.67-3.83 |
| Isoform 7 | Haplotype 7 | 4 | 2.71 | 2.33-3.00 |
| Isoform 8 | Haplotype 8 | 3 | 3.50 | 3.33-3.67 |
| Isoform 9 | Haplotype 9 | 3 | 2.72 | 1.83-3.17 |
| Isoform 10 | Haplotype 11 | 1 | 2.50 | - |
| Isoform 11 | Haplotype 13 | 1 | 2.50 | - |
| Isoform 12 | Haplotype 14 | 1 | 3.17 | - |
| Isoform 13 | Haplotype 15 | 1 | 2.17 | - |
| Isoform 14 | Haplotype 16 | 1 | 2.17 | - |
| Isoform 15 | Haplotype 17<sup>α</sup> | 1 | 0.17 | - |
| Isoform 16 | Haplotype 18<sup>α</sup> | 1 | 1.17 | - |
| Isoform 17 | Haplotype 19<sup>α</sup> | 1 | 0.17 | - |
| Isoform 18 | Haplotype 20<sup>α</sup> | 1 | 0.50 | - |
| Isoform 19 | Haplotype 21<sup>α</sup> | 4 | 0.36 | 0.00-0.50 |
| Isoform 20 | Haplotype 22<sup>α</sup>, 28<sup>αδ</sup> | 4 | 0.72 | 0.50-1.00 |
| Isoform 21 | Haplotype 23<sup>δ</sup> | 1 | 2.50 | - |
| Isoform 22 | Haplotype 24<sup>δ</sup> | 1 | 2.67 | - |
| Isoform 23 | Haplotype 25<sup>δ</sup> | 1 | 3.17 | - |
| Isoform 24 | Haplotype 26<sup>δ</sup> | 1 | 1.00 | - |
| Isoform 25 | Haplotype 27<sup>δ</sup> | 1 | 3.67 | - |
| Isoform 26 | Haplotype 29<sup>δ</sup> | 1 | 2.67 | - |
| Isoform 27 | Haplotype 30<sup>δ</sup> | 1 | 1.50 | - |
| Isoform 28 | Haplotype 31<sup>δ</sup> | 1 | 2.17 | - |
| Isoform 29 | Haplotype 32<sup>δ</sup> | 1 | 3.33 | - |
| Isoform 30 | Haplotype 33<sup>δ</sup> | 1 | 2.67 | - |
| Isoform 31 | Haplotype 34<sup>δ</sup> | 1 | 3.50 | - |
| Isoform 32 | Haplotype 35<sup>δ</sup> | 1 | 3.17 | - |
| Isoform 33 | Haplotype 36<sup>δ</sup> | 1 | 3.33 | - |
| Isoform 34 | Haplotype 37<sup>δ</sup> | 1 | 3.50 | - |
| Isoform 35 | Haplotype 38<sup>δ</sup> | 1 | 3.17 | - |
| Isoform 36 | Haplotype 39<sup>δ</sup> | 1 | 3.83 | - |

<sup>α</sup>Haplotypes that contain a premature stop codon for *SnTox5*. <sup>δ</sup>Haplotypes with a sequence coverage between 50 to 100% for *SnTox5*,
□ Analysis of variance of average disease reaction was performed only for the isoforms represented by more than ten isolates. Least significant difference was calculated at the 0.05 level of probability. Average disease reactions followed by same letter are not significantly different at the 0.05 level probability.

The Upper Midwest population, which included the reference isolate Sn2000, consisted of five haplotypes defined by six non-synonymous and two synonymous polymorphisms (Table 2). A nucleotide diversity (Pi) of 0.00109 was observed in the Upper Midwest population, which was lower than that of the South/East population. The South/East population consisted of nine haplotypes defined by seven non-synonymous and one synonymous polymorphism. The nucleotide diversity within the South/East population was calculated at 0.00327, higher than that of the Upper Midwest population.

The calculated pN/pS ratio for the Upper Midwest population was 0.95 (Table 2), however the calculated pN/pS ratio for the South/East population was substantially higher at 2.22 (Table 2), suggesting that SnTox5 was undergoing purifying selection in the Upper Midwest but strong diversifying selection in the South/East. Calculation of the pN/pS ratios only for the mature protein coding region of the gene gave similar results with pN/pS ratios of 0.47 and 1.89 for the Upper Midwest and South/East population respectively (Table 2). The pN/pS ratios were not calculated for the Oklahoma population due to a complete lack of synonymous SNPs. However, in the Oklahoma population, six nonsynonymous mutations were identified in the complete gene with five of these affecting the mature protein indicating diversifying selection pressure in this region. The Oregon population was made up of a single haplotype and therefore no calculation was possible. These results indicated that SnTox5 was undergoing different types of selection in different regions of the United States, likely adapting to the locally planted cultivars.

### SnTox5 isoform variation contributes to quantitative levels of virulence

The 22 haplotypes encoded 20 different isoforms of SnTox5 (Supplementary Figure 3). Three active and five inactive isoforms of SnTox5 were identified in the Upper Midwest population (n=56), where isoform 1 was the most prevalent at 60.71% (Figure 3). Furthermore, 94.44% of the isolates that harbored isoform 1 were from the Upper Midwest population. The Southern/Eastern population (n=48) harbored nine isoforms (eight active and one inactive) with isoform 2 and 3 representing 35.42% and 33.33% of the population, respectively (Figure 3). In addition, 70.83% of the isolates that contained isoform 2 and 100% of the isolates that contained isoform 3 were from the South/East population (Figure 3). The *P. nodorum* population from Oklahoma (n=17) harbored eight isoforms (Figure 3). Isoform 5 was the most prevalent form of the Oklahoma population, consisting of 35.29% of the population. The Oregon population consisted of only seven isolates where all of them harbored isoform 2 (Figure 3). No isoform was identified in all four of the populations. Like the haplotype analysis, the isoform analysis shows a higher diversity in SnTox5 in the South/East and Oklahoma populations compared to the Upper Midwest population.

**Figure 3.**
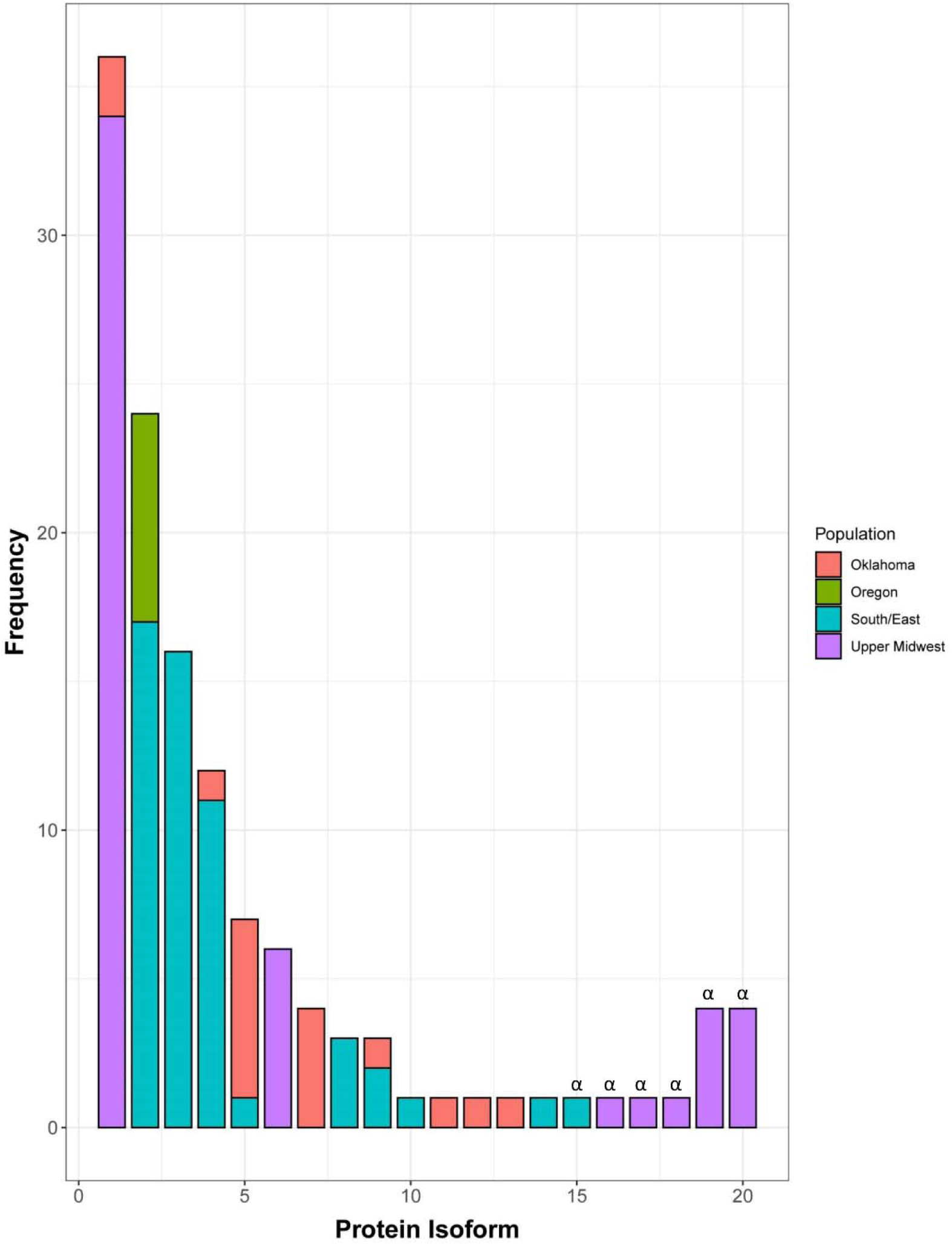
Prevalence of isoforms of SnTox5 in Upper Midwest, South/East, Oregon and Oklahoma population of *P. nodorum*. Distribution of isoforms of SnTox5 in different *P. nodorum* populations from four regions of the United States showed that each population consisted of multiple isoforms of SnTox5, except the Oregon population, with varying degree of prevalence. SnTox5 isoform1 was the most prevalent isoform of the Upper Midwest population whereas isoform 5 was the most prevalent in the Oklahoma population. Isoform 2 was the most prevalent in both the Oregon and South/East populations. Isoforms marked with ‘α’ represent an inactive form of SnTox5 with a premature stop codon.

To examine the likely effect of non-synonymous substitutions on virulence, SnTox5 isoforms produced by greater than 10 individuals were statistically analyzed based on their average disease reaction on LP29. Therefore, isoforms 1, 2, 3, and 4 that were represented by 36, 24, 16 and 12 isolates, respectively, were used. The reference isolate Sn2000 produced isoform 1 and was included in the 36 isolates with isoform 1 for this analysis. Isolates expressing isoform 1 (produced by 34 isolates from the Upper Midwest and two isolates from Oklahoma) and isoform 4 (produced by five isolates from the Upper Midwest, one isolate from Oklahoma and six isolates from South/East population) caused average disease reactions of 2.27 and 2.23, respectively on LP29, and were not significantly different from each other (Table 3). The isolates producing isoform 2 (produced by seven isolates from Oregon and 17 isolates from the South/East population) and the isolates producing isoform 3 (produced by 16 isolates only from the South/East population) showed an average disease reaction of 2.96 and 3.36, respectively, on LP29. The average disease reactions of isolates producing isoform 3 were significantly higher than those producing isoform 2 at the 0.05 level of probability (Table 3). Average disease reactions of isolates harboring both isoform 2 and 3, had significantly higher average disease reactions than those of isoforms 1 and 4 at the 0.05 level of probability (Table 3) indicating that variation in the SnTox5 amino acid complement influenced the level of virulence of the isolates producing them, likely due to the direct or indirect interaction with Snn5. Analysis of the amino acid sequence of each isoform showed that isoform 2 had a T155R substitution and isoform 3 had a T155K substitution compared to isoforms 1 and 4 suggesting that the amino acid at the 155^th^ position contributed significantly to the variation in average disease reaction (Figure 4).

**Figure 4.**
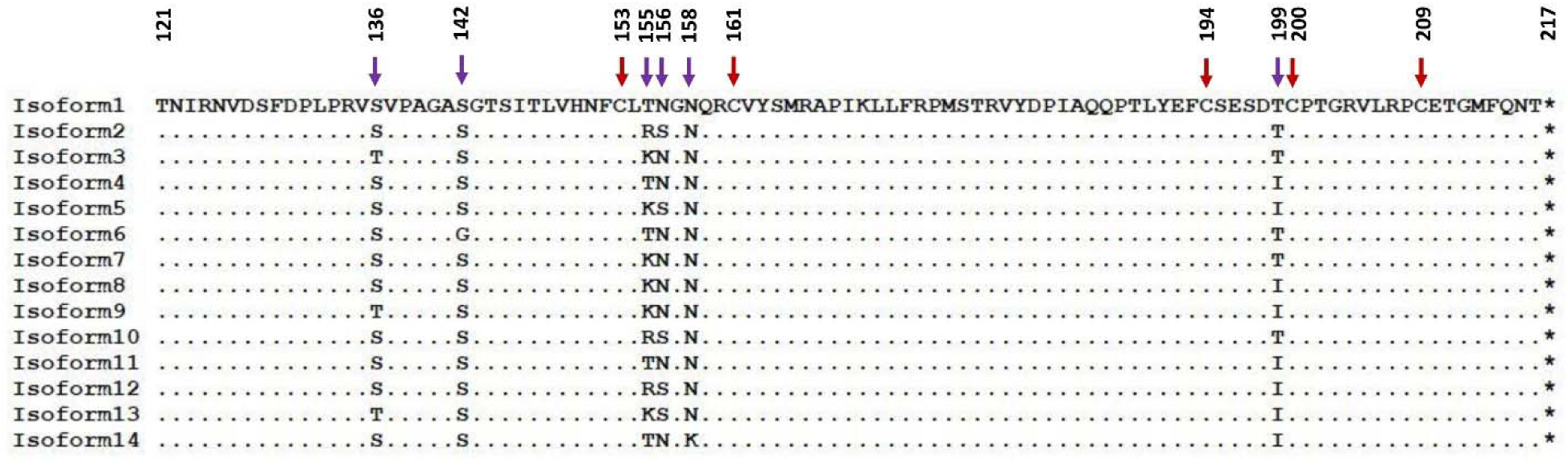
A portion of the amino acid sequence of the active isoforms of SnTox5 representing the critical substitutions that contribute to the variation in disease using isoform 1 as a reference. Purple arrows indicate the physical position of the substitutions and red arrows indicate the physical position of the cysteine residues. The 155^th^ position consisted of either threonine (T), arginine (R), or lysine (K) whereas the 156^th^ position consisted of either asparagine (N) or serine (S) and was flanked by two cysteines at the 153^rd^ and 161^st^ positions which were predicted to form a di-sulfide bond. Substitutions T155R and T155K contributed to an increase in averaged disease reaction type whereas N156S contributed to the variation in average disease reaction type on LP29.

When comparing all 14 active SnTox5 isoforms, amino acid substitutions were frequently observed at the 155^th^ and 156^th^ amino acid positions (Figure 4). Three amino acids including threonine (T), arginine (R), and lysine (K), were observed at the 155^th^ position and two amino acids including asparagine (N) and serine (S) were observed at the 156^th^ position (Table 4). Isolates of the 14 isoforms were assembled into groups based on the amino acids at the 155^th^ and 156^th^ positions and amino acid combinations of T-N, R-S, K-S, and K-N showed average disease reactions of 2.31, 2.95, 2.95 and 3.20, respectively (Table 4). Isolates with R-S, K-S, and K-N substitutions caused significantly higher average disease reaction compared to the T-N substitution, suggesting that variation in amino acid residues at these two positions contributed significantly to the virulence of the isolates producing them.

**Table 4.** Amino Acids substitutions at the 155^th^ and 156^th^ positions contribute to the variation in average disease reaction on LP29caused by the *P. nodorum* isolates harboring active isoforms of SnTox5.

| Amino acid at the position |  | Isoforms represented§ | Number of isolates with the substitution | Average disease reaction type□ |
| --- | --- | --- | --- | --- |
| 155<sup>th</sup> | 156<sup>th</sup> |  |  |  |
| T | N | 1, 4, 6, 11, 15 | 56 | 2.31a |
| R | S | 2, 10, 12 | 26 | 2.95b |
| K | S | 5,13 | 8 | 2.95b |
| K | N | 3,7,8,9 | 26 | 3.20b |

§ Isolates from these isoforms that represent identical substitutions were pooled together for the mean comparison of average disease reaction.
□ Least significant difference (LSD) was calculated at $P < 0.05$ probability. Numbers followed by the same letter were not significantly different at the 0.05 level of probability.

SnTox5 isoforms with a premature stop codon showed average disease reactions ranging from 0-1.17 on LP29 (Table 3) showing that isolates with a truncated SnTox5 failed to cause disease on LP29. These results indicated that SnTox5 may be evolving to become a more effective protein, increasing the virulence of the pathogen.

### SnTox5 expression peaks after penetration but prior to visible lesions

To examine the expression profile of *SnTox5* throughout the infection cycle, RT-qPCR was performed using *in-planta* samples. Expression of *SnTox5* in Sn2000 was analyzed at 4, 12, 24, 48, 72, 96, and 120 hpi on the durum wheat cultivar Lebsock. *SnTox5* expression peaked at 24 hpi, prior to the onset of lesions that typically became visible between 48 and 72 hpi (Figure 5). At 24 hpi, *SnTox5* was expressed approximately six times that of the *actin* gene. The expression of *SnTox5* gradually decreased with the progression of the disease before it returned to levels like that of *actin* at 120 hpi where the pathogen had already colonized the mesophyll tissue (Figure 5).

**Figure 5.**
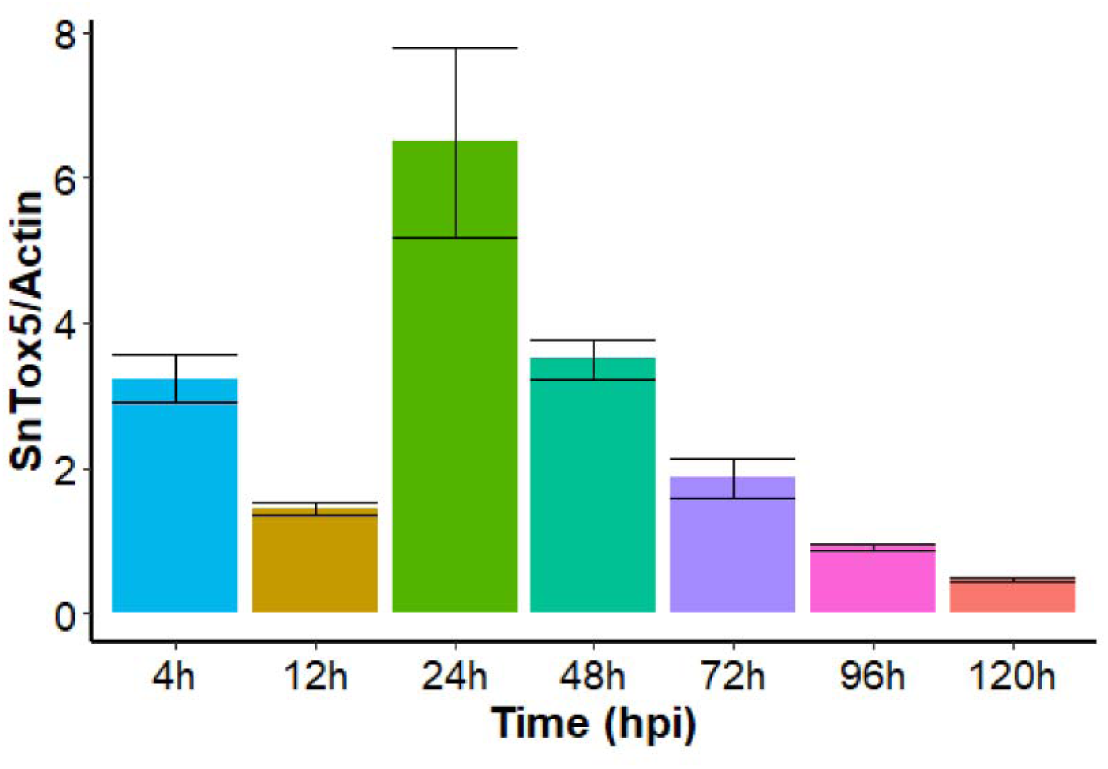
Temporal expression pattern of *SnTox5 in planta*, on Lebsock inoculated with Sn2000. The x-axis shows the sample collection time points for qPCR and the y-axis represents the expression of *SnTox5* relative to the expression of the *actin* gene. Error bars represent the standard error of the mean from three replications for each time point.

### Laser confocal microscopy shows that SnTox5 facilitates complete colonization of LP29

To visualize the effect of the SnTox5-*Snn5* interaction on penetration and colonization of the leaf cell layers, fluorescently labeled *P. nodorum* strains Sn2000, Sn2kΔTox5-15, Sn79+Tox5*-*3 and Sn79-1087 were inoculated onto LP29. The infection process of each strain was observed using laser scanning confocal microscopy at seven different time points post-inoculation. Germination of spores of all four strains was visible within 4 hpi (Figure 6 and Figure 7). At 12 hpi, penetration of the leaf surface was also visible for all four strains, however, it was clear that strains that contained *SnTox5* had increased penetration compared to strains that lacked *SnTox5* (Figure 6 and Figure 7).

**Figure 6.**
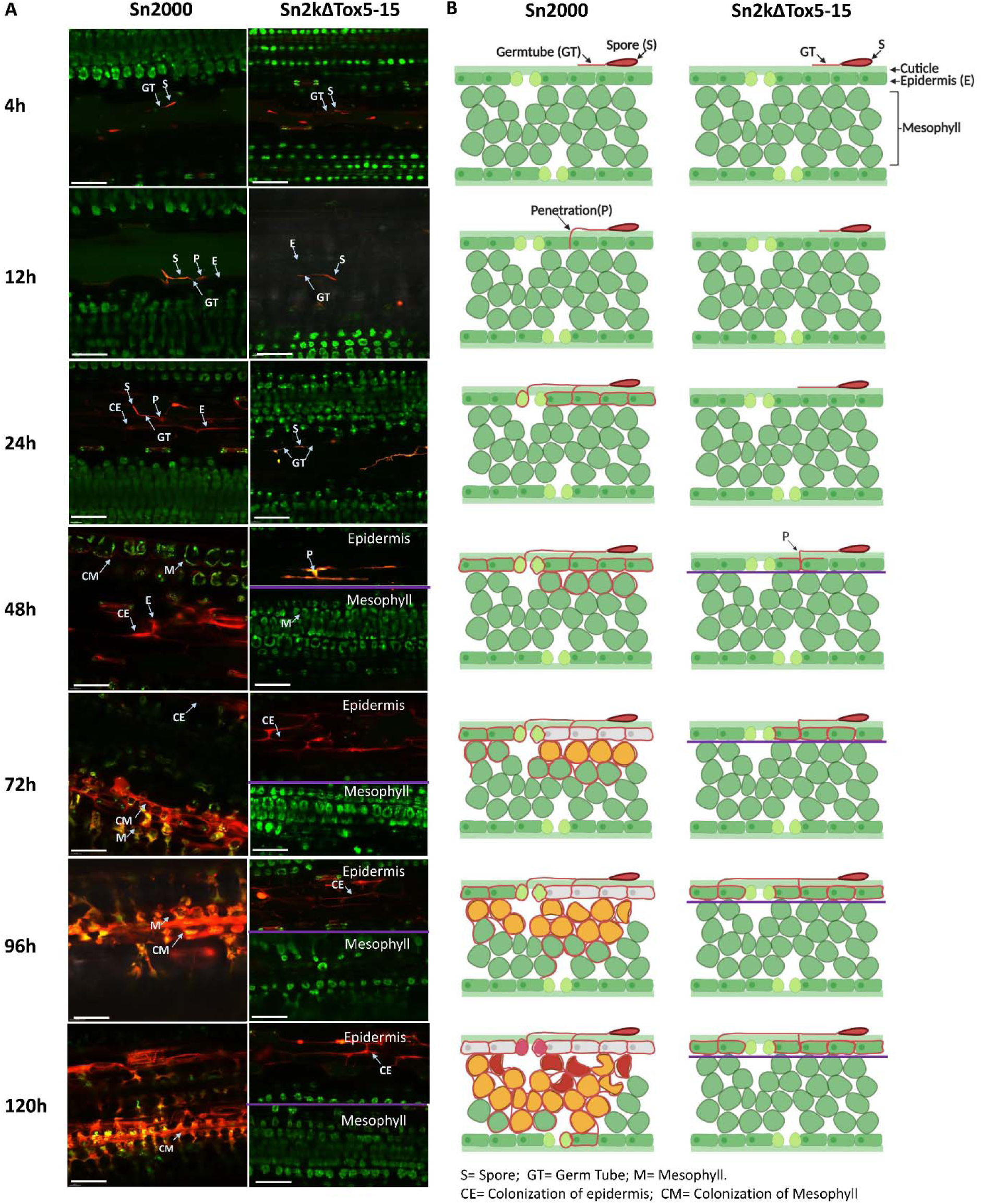
Laser confocal microscopy of the infection process of red fluorescent protein (RFP) tagged P. nodorum strains Sn2000 and Sn2kΔTox5-15 on wheat differential line LP29 (Snn5). (A) Micrographs of wheat leaves obtained through confocal imaging at 4, 12, 24, 48, 72, 96, and 120 hours post inoculation (hpi) of Sn2000 and Sn2kΔTox5-15. Fungal spores and hyphae are displayed in red. Wheat cells are displayed in various colors depending on the autofluorescence emitted by the degrading chloroplast, where green color indicates healthy cells and yellow to red color indicates cells that are undergoing programmed cell death. Separate z-stack micrographs taken at the epidermis and the mesophyll tissue at 48 to 120 hpi timepoints showed that Sn2kΔTox5-15 failed to advance into the mesophyll tissue. (B) Schematic drawings of transverse sections of each micrograph of (A). Purple line separates the epidermis from the mesophyll tissue. Scale bar = 60 µm.

**Figure 7.**
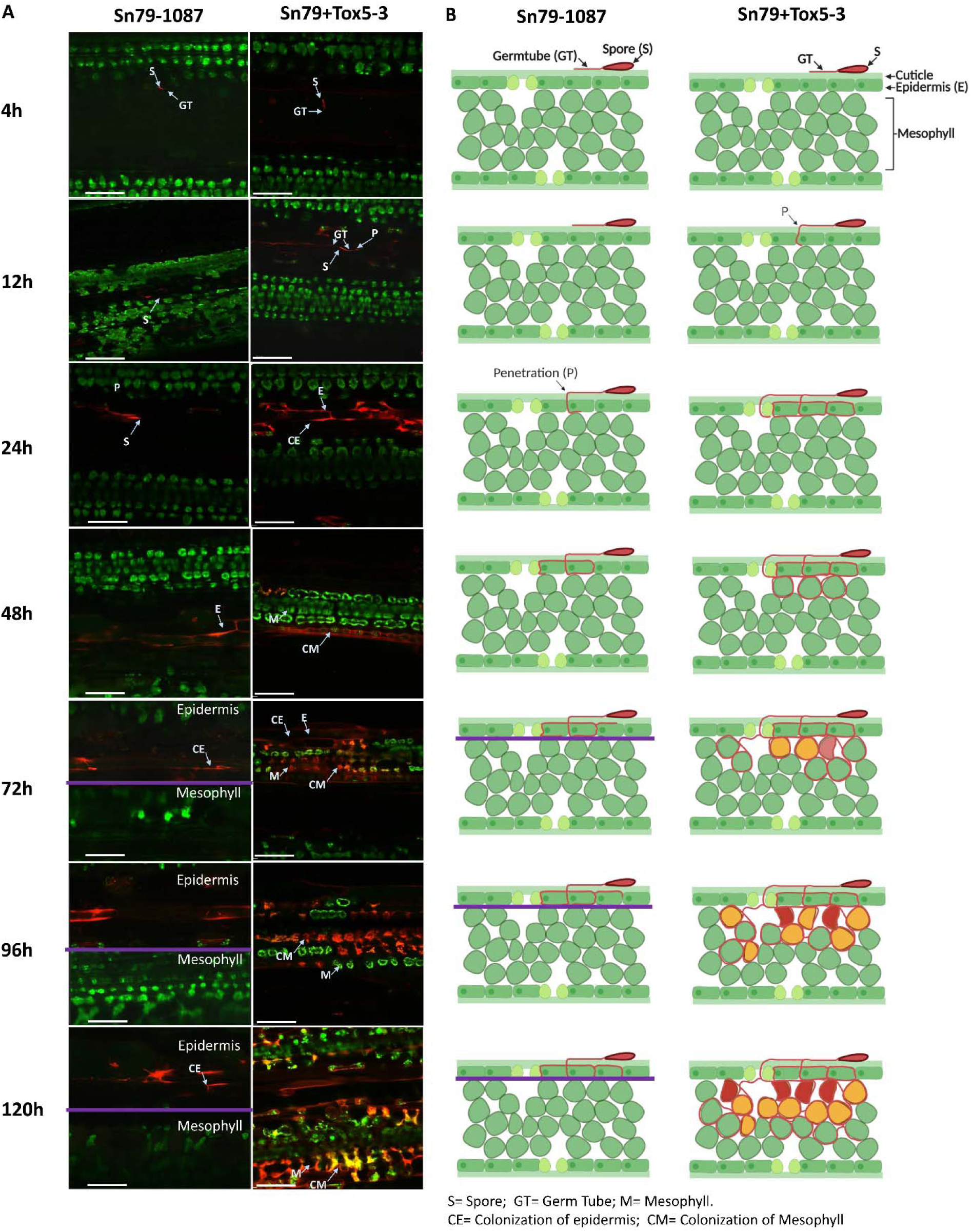
Laser confocal microscopy of the infection process of red fluorescent protein (RFP) labeled *P. nodorum* strains Sn79-1087 (avirulent) and Sn79+Tox5-3 on wheat differential line LP29 (*Snn5*). (A) Micrographs of wheat leaves obtained through confocal imaging at 4, 12, 24, 48, 72, 96, and 120 hours post inoculation (hpi) of Sn79-1087 and Sn79+Tox5-3. Fungal spores and hyphae are displayed in red. Wheat cells are displayed in various colors depending on the auto florescence emitted by the degrading chloroplast, where green color indicates healthy cells and yellow to red color indicates cells that undergoing program cell death. Separate z-stack micrographs taken at the epidermis and the mesophyll tissue at 72 hpi to 120 hpi timepoints showed that Sn79-1087 failed to reach the mesophyll tissue as *SnTox5* mutant of Sn2000. Transfer of *SnTox5* to Sn79-1087 enables the fungus to reach the mesophyll tissue. (B) Schematic drawings of transverse sections of each micrograph of (A). Purple line separates the epidermis from the mesophyll tissue. Scale bar = 60 µm.

At 48 hpi, all strains were able to colonize the epidermal tissue and Sn2000 and Sn79+Tox5-3 had initiated colonization of the vascular and mesophyll tissue, however, neither Sn2kΔTox5-15 nor Sn79-1087 were ever able move past the epidermal layer (Figure 6 and Figure 7). For Sn2000 and Sn79+Tox5-3 the breakdown of chloroplasts in the mesophyll cells surrounded by the fungal hyphae was observed at 72 hpi, turning chloroplasts from green to yellow (Figure 6 and Figure 7). Deterioration of the chloroplasts was followed by the shrinking and total collapse of the surrounded mesophyll cells at the 96 and 120 hpi time points (Figure 6 and Figure 7). By 120 hpi, the majority of the mesophyll tissue was colonized by both Sn2000 and Sn79+Tox5-3. It was evident that Sn2000 colonized more tissue than that of Sn79+Tox5-3 at 120 hpi, indicating that additional virulence factors effective on wheat may be present in Sn2000 that are not present in the avirulent Sn79-1087 (Figure 6 and Figure 7).

Even though Sn2kΔTox5-15 and Sn79-1087 lacked *SnTox5*, both penetrated the epidermis by 24 hpi. Extensive colonization of the epidermis was observed by 96 hpi, similar to Sn2000 and Sn79+Tox5-3. However, unlike Sn2000 and Sn79+Tox5-3, Sn2kΔTox5-15 and Sn79-1087 were unable to colonize the mesophyll tissue or the vascular tissue of the leaf (Figure 6 and Figure 7).

Typically, lesions started to appear on the leaves between 48 and 72 hpi, correlating with the colonization of the mesophyll tissue. Until *P. nodorum* started to colonize the mesophyll tissue, the leaf remained green. Both Sn2kΔTox5-15 and Sn79-1087 failed to produce lesions on inoculated leaves. In contrast, Sn2000 and Sn79+Tox5-3 were able to reach the mesophyll and vascular tissue by 48-72 hpi, coinciding with the emergence of visible leaf necrosis.

As a proxy for fungal fitness, the fungal volume of Sn2000 and Sn2kΔTox5-15 were measured at 12, 24, 48, 72, 96, and 120 hpi (Figure 8A). The volume of Sn2000 gradually increased on LP29 over time as expected, however, the volume of Sn2kΔTox5-15 on LP29 remained constant from 12 to 120 hpi. The volume of Sn2000 started to increase significantly at 24 hpi, coinciding with the upregulation of *SnTox5* and fungal colonization of the mesophyll layer (Figure 6, 8A). Collectively, the microscopic analysis of leaf penetration and colonization showed that SnTox5 clearly facilitates colonization of the mesophyll layer of the leaf, followed by PCD through its targeting of the *Snn5* pathway, providing nutrient for the completion of its pathogenic life cycle.

**Figure 8.**
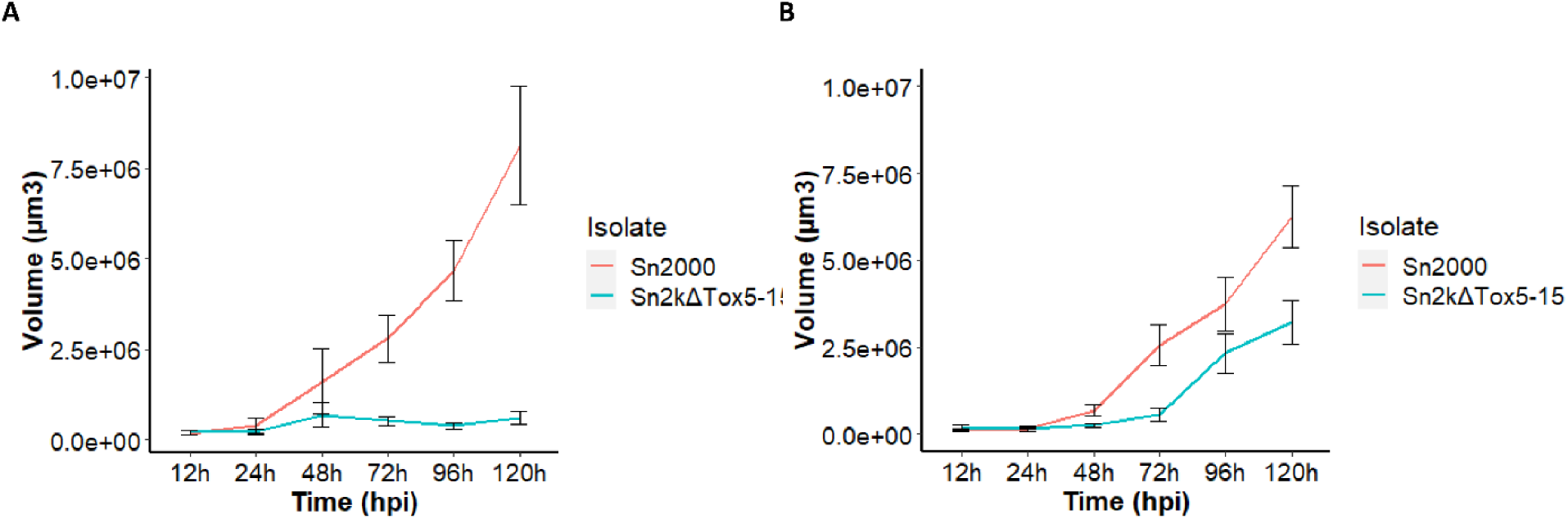
Volume of the fungus calculated through laser confocal microscopy on A). LP29 (*Snn5*) and B). Lebsock (*Snn5* and *Tsn1*) for the strains Sn2000 (+*SnTox5*, +*SnToxA*) and Sn2kΔTox5(-SnTox5, +SnToxA) at various time points post inoculation. The x-axis represents hours post inoculation (hpi) and the y-axis represents the volume of the fungus in µm^3^. The volume of Sn2000 increased linearly after 24 hpi on both wheat lines. The increase in volume of Sn2kΔTox5 on LP29 was negligible during the experiment. However, linear increase in volume of Sn2kΔTox5 was observed after 72 hpi on Lebsock due to the establishment of SnToxA-Tsn1 interaction.

### SnTox5 facilitates colonization of the epidermal layer of LP29 even in the absence of Snn5

To further analyze the function of the *SnTox5-Snn5* interaction, we inoculated Sn2000 and the Sn2kΔTox5-15 mutant on LP29 and its *Snn5* disruption mutant LP29Δsnn5. At 120 hpi, Sn2000 was able to colonize both the epidermis and the mesophyll tissue as described above (Figure 9). In the Sn2000 inoculation of LP29Δsnn5, Sn2000 was able to progress into the mesophyll tissue, however, unlike the Sn2000 - LP29 inoculation, the fungus was not able to induce PCD and was only observed to advance into the first two cell layers of the mesophyll tissue. As mentioned above, *SnTox5* mutants of Sn2000 were only able to penetrate and colonize the epidermal tissue of LP29. At 120 hpi, spores of Sn2kΔ Tox5-15 were able to form germ tubes on LP29Δsnn5. Unexpectedly, Sn2kΔTox5-15 was not even able to penetrate the epidermal tissue of LP29Δsnn5, even up to 120 hpi (Figure 9). More work will need to be done on this interaction to understand the role of *Snn5* in penetration in the absence of SnTox5.

**Figure 9.**
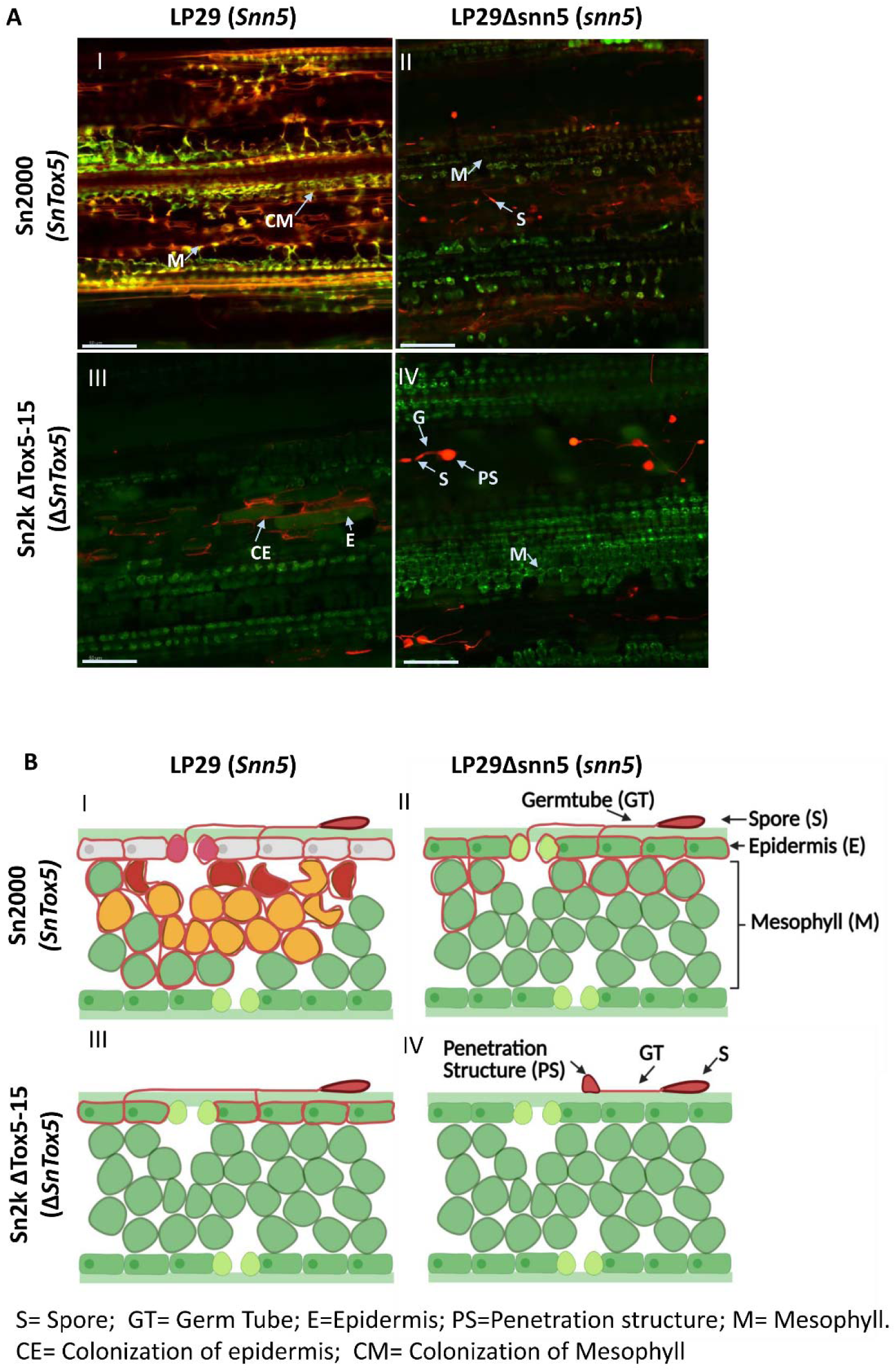
Laser confocal microscopy of red fluorescent protein (RFP) tagged Sn2000 and Sn2kΔTox5-15 inoculated on LP29 and LP29Δsnn5 at 120 hpi. (A). Micrographs of RFP tagged Sn2000 (+*SnTox5*) and Sn2kΔTox5 (-SnTox5) inoculated on LP29 (Snn5), and LP29Δsnn5 (*snn5*) at 120 hpi. (B) Schematic drawings of transverse sections of each micrograph of (A). A(I) and B(I). Sn2000 induced programmed cell death (PCD) and colonized the mesophyll tissue of LP29. A(II) and B(II). Sn2000 was able to colonize the epidermis and hyphae were advanced into the mesophyll tissue but failed to induce PCD on LP29Δsnn5 due to the lack of functional Snn5. A(III) and B(III). Sn2kΔTox5 colonized the epidermal tissue but failed to progress to the mesophyll tissue since it lacked *SnTox5*. A(IV) and B(IV). Sn2kΔTox5 formed a penetration structure on LP29Δsnn5. However, Sn2kΔTox5 was not able to penetrate the epidermis since Sn2000ΔTox5 and LP29Δsnn5 lacked *SnTox5* and *Snn5*. Therefore, these results showed establishment of SnTox5-Snn5 is essential for the *P. nodorum* to colonize the mesophyll of LP29 and lack of either partner of the interaction cause a deleterious effect of fungal growth. Scale bar = 100 µm.

### Laser confocal analysis of Sn2000 and Sn2kΔTox5-15 on Lebsock

To further evaluate the additive nature of SnTox5-*Snn5* and SnToxA-*Tsn1* interactions, we analyzed the infection process of Sn2000 and Sn2kΔTox5-15 on Lebsock, which carries both *Tsn1* and *Snn5*. Both Sn2000 and Sn2kΔTox5-15 were able to penetrate and colonize the epidermis of the wheat line Lebsock within 24 hpi (Figure 10). Sn2000 was able to penetrate the mesophyll layer by 48 hpi (Figure 10). However, Sn2kΔTox5-15 did not reach the mesophyll tissue until 96 hpi (Figure 10) showing that the SnTox5-*Snn5* interaction was facilitating a more rapid colonization of the mesophyll tissue than the SnToxA-*Tsn1* interaction alone.

Calculation of the fungal volume of Sn2000 and Sn2kΔTox5-15 in Lebsock, clearly showed that even though both strains were able to colonize the mesophyll tissue, Sn2000 had significantly higher fungal volume compared to that of the Sn2kΔTox5-15, starting at 48 hpi and continuing through the 120 hpi time point (Figure 8B). In addition, calculation of fungal volume at each time point showed that the rate of the increase in fungal volume was present at 24 hpi for Sn2000, whereas the fungal volume did not increase until 72 hpi for Sn2kΔTox5-15 (Figure 8B). These results suggest once again, SnTox5 is facilitating entry into the mesophyll and that establishment of one NE-sensitivity gene interaction is sufficient to colonize the mesophyll layer, but the two interactions act synergistically during the colonization process to increase pathogen fitness and therefore, the rate of colonization.

**Figure 10.**
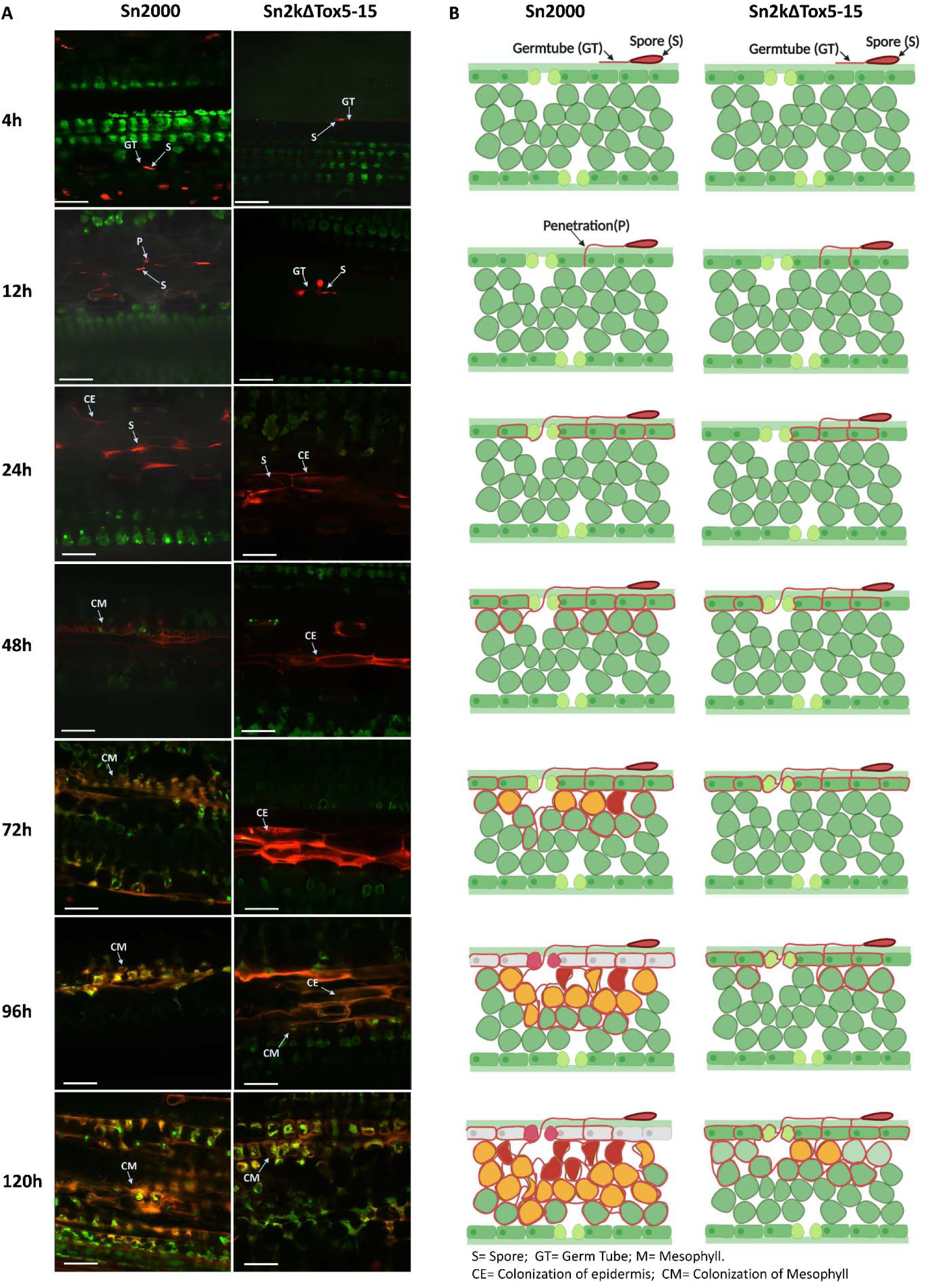
Laser confocal microscopy of the infection process of red fluorescent protein (RFP) labeled *P. nodorum* strains Sn2000 and Sn2kΔTox5-15 on wheat cultivar Lebsock (*Snn5* and *Tsn1*). (A) Micrographs of wheat leaves obtained through laser confocal imaging at 4, 12, 24, 48, 72, 96, and 120 hpi of Sn2000 and Sn2kΔTox5. Fungal spores and hyphae are displayed in red. Wheat cells are displayed in various colors depending on the autofluorescence emitted by the degrading chloroplast, where green color indicates healthy cells and yellow to red color indicates cells that are undergoing programmed cell death. (B) Schematic drawings of transverse sections of each micrograph of (A) clearly illustrate that Sn2000 was able to advance into the mesophyll tissue and colonize the mesophyll tissue rapidly compared to the that of Sn2kΔTox5-15. Scale bar = 60 µm.

## Discussion

*P. nodorum* is a necrotrophic fungal pathogen that deploys a plethora of necrotrophic effectors (NEs) to induce programmed cell death (PCD) on susceptible wheat lines. In this study, we performed GWAS on a *P. nodorum* natural population of 197 isolates, to identify *Sn2000_06735* as a candidate for *SnTox5*. We subsequently used CRISPR/Cas9 mediated gene editing to disrupt the gene and gain-of -function transformation to complement the gene showing that *Sn2000_06735* was both sufficient and necessary to cause disease on LP29, the *Snn5* differential line.

Phenotyping, and subsequent QTL analysis on the LP749 population that segregated for sensitivity to SnTox5, showed that the disruption of *Sn2000_06735* eliminated any disease association with *Snn5*. Sn79-1087 also had no disease association with *Snn5*, however, strong disease association with *Snn5* was detected when inoculated with the Sn79-1087 strain that was transformed with *Sn2000_06735*. Therefore, GWAS, gene disruption, gain-of-function transformation, and QTL analysis on the LP749 wheat population validated that *Sn2000_06735* was *SnTox5*.

*SnTox5* encodes an immature 217 amino acid small secreted protein that harbors a 22 amino acid secretion signal and a putative 45 amino acid pro-sequence that is cleaved at a predicted Kex2 protease cleavage site. BLASTp search with the SnTox5 amino acid sequence showed that SnTox3 was the only hit in the NCBI non-redundant database and only showed 45% homology. Although this is not a high level of homology, it is interesting that SnTox3 is also a pre-pro protein with a Kex2 cleavage site (Liu et al. 2009; Outram et al. 2020) and structural homology is also observed based on similar placement of the cysteine residues, indicating a potentially similar mature protein structure. The Kex2 protease is unique to fungi and Outram et al. (2020) showed that Kex2-processed pro-domain (K2PP) effectors were common in pathogenic fungi and included the *P. nodorum* effectors SnTox3 and SnToxA and now SnTox5. Outram et al. (2020) also presented the crystal structure of SnTox3 and experimentally demonstrated that the Kex2-processed pro-domain was critical for SnTox3 folding and activity. The validation of SnTox5 provides another K2PP effector interaction for further study of this class of effectors.

Among the *P. nodorum* isolates used in this study, 75.6% harbored a functional *SnTox5* gene. The level of prevalence observed for *SnTox5* was only slightly higher than that of *SnToxA* and *SnTox3*, which were found in 63.4% and 58.9% of the isolates in the same collection used in this study, respectively (Richards et al. 2019). In contrast, the presence of *SnTox5* was less compared to the prevalence of *SnTox1* and *SnTox267* in the same population, which was 95.4% for each (Richards et al. 2019; Richards et al. 2021). The ability to target multiple host susceptibility genes such as in SnTox267 (Richards et al. 2021) or SnTox3 (Friesen et al. 2008; Zhang et al. 2011; Zhang et al. 2021), the existence of a secondary function such as the ability of SnTox1 to bind chitin (Liu et al. 2016) or SnTox3 to target PR1 proteins (Sung et al. 2020), or the prevalence of host susceptibility genes in the planted wheat of a given region (Richards et al. 2019), all likely govern the frequency of an effector gene in a fungal population. Therefore, prevalence of *SnTox5* in the majority of *P. nodorum* isolates collected from the Upper Midwest, Oklahoma, Oregon and South/East regions of US suggests the prevalence of *Snn5* in wheat planted in these regions or an additional function that drives the maintenance of this gene.

Richards et al. (2019) used the same US *P. nodorum* population to show that genes predicted to encode effectors were under stronger diversifying selection compared to genes encoding for secreted non-effectors or non-secreted proteins. Richards et al. (2019) also compared the Upper Midwest and South/East populations to show that several effector genes were under diversifying selection in one population but under purifying selection in the other population(s). One of these genes was SnTox3 which was under purifying selection in the South/East population and diversifying selection in the Upper Midwest population. Here we show that SnTox5 is under purifying selection in the Upper Midwest but diversifying selection in the South/East population, the opposite of SnTox3. Because the Upper Midwest wheat region is predominately spring wheat and the South/East wheat region is predominately winter wheat, the locally planted cultivars are vastly different. These differences include the complement of effector targets (e.g. *Snn3* for SnTox3 and *Snn5* for SnTox5) present in the local cultivars (Crook et al. 2012). Additionally, it is likely that population specific alleles of *Snn3* and *Snn5* exist that may be driving the diversification of both SnTox3 and SnTox5 in their respective populations.

*P. nodorum* isolates harboring a diversity of active isoforms of the SnTox5 protein were evaluated for disease reaction on the *Snn5* differential line LP29. Isolates harboring SnTox5 isoforms predominately from the South/East winter wheat growing regions were significantly more virulent on LP29 than isolates harboring isoforms that were prevalent in the Upper Midwest spring wheat and durum wheat regions. No isolates producing the most virulent two isoforms were identified in the Upper Midwest population indicating that the genetic background of the winter wheat *P. nodorum* population of the South/East is likely the selection pressure driving this diversity and the increased virulence.

The importance of critical amino acid residues in *P. nodorum*-wheat effector-target gene interactions have been reported for SnToxA (Meinhardt et al. 2002; Lu et al. 2014) and SnTox3 (Sung et al. 2021). A total of 14 active isoforms of SnTox5 were identified in our natural population. Of the four major isoforms, isolates carrying isoforms with T155K and T155R amino acid substitutions caused significantly higher disease. Threonine (T) is neutral in its hydrophobicity whereas both lysine (K) and arginine (R) are highly hydrophilic residues. Therefore, increase in the hydrophilicity at the 155^th^ position appears to result in an increase in virulence of the isolates producing these isoforms. Amino acid residues at the 156^th^ position were also variable and consisted of either serine (S) or asparagine (N). The highest average disease reaction was observed when position 155 and 156 were occupied by K and N compared to K and S, respectively, however, these differences were not significant at a 0.05 level of probability. Based on our results, we hypothesize that the amino acid residues at position 155 and possibly 156 are under selection and are critical to the effectiveness of the protein in the SnTox5-*Snn5* interaction.

SnTox5 expression peaked early with its highest expression at 24 hpi with a gradual decrease through 120 hpi (Figure 5). This expression pattern indicated that SnTox5 was likely involved in the early colonization of the leaf including the initial colonization of the mesophyll, which is initiated at 24 hpi and continues through 120 hpi (Figure 6).

Laser scanning confocal microscopy was then used to evaluate the importance of SnTox5 in the various stages of infection. At 48 hpi, all strains with or without *SnTox5* were able to colonize the epidermal layer, indicating that SnTox5 was not necessary to colonize the epidermal layer of LP29. The visible differences in colonization began at 48 hpi where the strains producing SnTox5 were able to begin colonizing the mesophyll layer but those strains not producing SnTox5 were not.

To further investigate the function of SnTox5 in the presence and absence of *Snn5*, four combinations were evaluated using laser confocal microscopy including 1) Sn2000 (SnTox5) on LP29 (Snn5), 2) Sn2000 (SnTox5) on LP29Δsnn5 (no Snn5), 3) Sn2kΔTox5 (no SnTox5) on LP29 (Snn5) and 4) Sn2kΔTox5 (no SnTox5) on LP29Δsnn5 (no Snn5). In combination 1 the presence of *Snn5* and the production of SnTox5 resulted in full colonization of the epidermis and mesophyll and complete cellular breakdown by 120 hpi. In combination 2, where SnTox5 was produced but *Snn5* was absent, mycelium penetrated the epidermis as well as initiating the colonization of the first layers of the mesophyll, but mesophyll colonization was halted prematurely, likely due to the lack of PCD. In combination 3, where SnTox5 was not produced but *Snn5* was present, the pathogen was able to penetrate and colonize the epidermis, however, no colonization of the mesophyll was ever observed. In combination 4, an unexpected result was found. The combination of Sn2kΔTox5 and LP29Δsnn5 eliminated both the production of SnTox5 and the *Snn5* host gene. Our expectation was that the Sn2kΔTox5 strain would penetrate like combination 3 due to the lack of SnTox5, however, no penetration was observed in any of the leaves examined. This result was puzzling, so it was repeated several times with the same result each time. Our only explanation for this phenomenon is that *Snn5* is somehow involved in communication with the pathogen but working out a model to explain this will require further work, including the cloning of *Snn5*. Combinations 1 through 3 along with the previous results on LP29, strongly suggested that SnTox5 is facilitating the colonization of the mesophyll layer even in the absence of the PCD induced by the SnTox5-*Snn5* interaction.

To evaluate the role of SnTox5 and *Snn5* in the presence of SnToxA and *Tsn1*, we used laser confocal microscopy to collect fungal volume data and to visualize the pathogen movement of *P. nodorum* isolate Sn2000 (SnToxA/SnTox5) and the *SnTox5*-disrupted mutant Sn2kΔTox5 (SnToxA only) on the durum wheat line Lebsock (*Tsn1*/*Snn5*) (Figure 10). Sn2000 behaved as presented previously where it began colonization of the mesophyll at 48 hpi with cell death beginning to be visible at 72 hpi and complete colonization and cellular disruption by 120 hpi. Sn2kΔTox5, which produces SnToxA but not SnTox5 could not breach the mesophyll layer at 48 or 72 hpi but did begin to colonize the mesophyll by 96 hpi with more advanced colonization by the 120 hpi timepoint. Although SnToxA does target the susceptibility gene *Tsn1* to induce PCD, having SnTox5 facilitates an earlier (by as much as 48 h) and stronger colonization of the mesophyll resulting in earlier PCD and therefore faster acquisition of cellular nutrients needed to complete the fungus’ life cycle. This combination reiterates that SnTox5 is facilitating mesophyll colonization, a role that is not replicated by SnToxA.

We are hypothesizing that SnTox5 has a secondary effector function that facilitates entry into the mesophyll that is important prior to its necrotrophic effector function that targets *Snn5*. This hypothesis is supported by results including the peak expression of SnTox5 at 24 hpi, a time point where the pathogen is initiating penetration into the mesophyll as well as the multiple pathogen strain – host genotype combinations that highlight the roles of *Snn5* and SnTox5. These include 1) *P. nodorum* isolate Sn2000 was able to enter the mesophyll of LP29 in the presence or absence of *Snn5*, however, SnTox5 nonproducing strains of these same isolates including Sn79-1087 and Sn2kΔTox5 were limited to the epidermis. 2) Inoculations of Sn2000 (SnToxA/SnTox5) and Sn2kΔTox5 (SnToxA alone) on the durum wheat cultivar Lebsock (*Tsn1/Snn5*) showed that Sn2000 penetrated and colonized both the epidermal and mesophyll layers by 48 hpi, however, Sn2kΔTox5 which only has *SnToxA* was not able to initiate colonization of the mesophyll layer until 96 hpi, indicating that SnTox5 was responsible for the rapid colonization of the mesophyll and that SnToxA was not nearly as efficient in this regard.

In this study we used GWAS with 197 *P. nodorum* isolates to identify a SnTox5 candidate gene followed by gain-of-function transformation and CRISPR-Cas9 based gene disruption for validation of the SnTox5 candidate. Using the same US population of *P. nodorum* isolates that collected from both winter and spring wheat regions of the US showed that the *SnTox5* gene was under purifying selection in the spring wheat growing region but under diversifying selection in the South/East US and Oklahoma winter wheat growing regions. One region of *SnTox5* was under strong diversifying selection and the two amino acid residues encoded by this region was contributing quantitatively to virulence. Additionally, we have shown multiple lines of evidence that SnTox5 is clearly facilitating early colonization of the mesophyll. Our current working model is that SnTox5 expression peaks early (24 hpi) where it facilitates the colonization of the mesophyll as early as 48 hpi, putting the pathogen in position to obtain nutrients that are a result of the SnTox5-*Snn5* induced PCD.

