## Supplemental Figure Legends for "The *Parastagonospora nodorum* necrotrophic effector SnTox5 targets the wheat gene *Snn5* and facilitates entry into the leaf mesophyll"

Supplementary Figure1. Nucleotide sequence of the Sn2000 SnTox5 gene used to transform Sn79-1087 to create the gain of function transformants Sn79+Tox5-3 and Sn79+Tox5-4 and the resulting amino acid sequence of SnTox5. Yellow color indicates the putative TATAA box 171 bp upstream of the start codon. Purple, red and green colors indicate the signal peptide, the pro-domain and the putative Kex2 site of the protein.

Supplementary Figure 2. Pairwise alignment of SnTox3 and SnTox5 amino acid sequence.

Supplementary Figure 3. Pairwise alignment of 20 isoforms using isoform 1 as the reference.
