## Supplementary figures and images for "The *Parastagonospora nodorum* necrotrophic effector SnTox5 targets the wheat gene *Snn5* and facilitates entry into the leaf mesophyll"

### Supplemental Figure 1

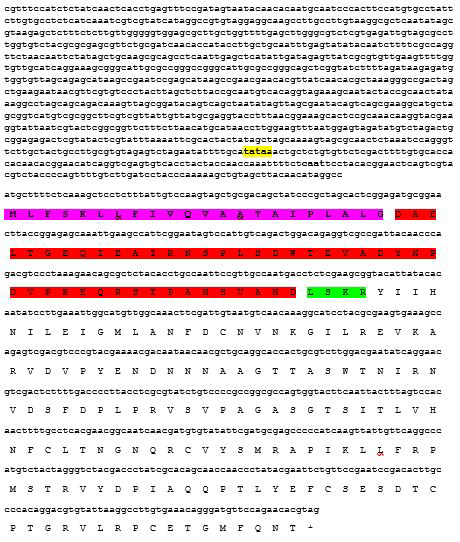

### Supplemental Figure 2

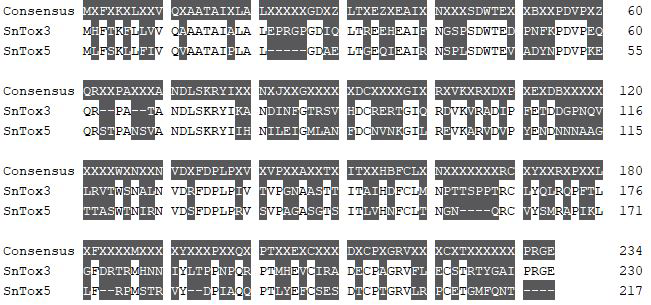

### Supplemental Figure 3

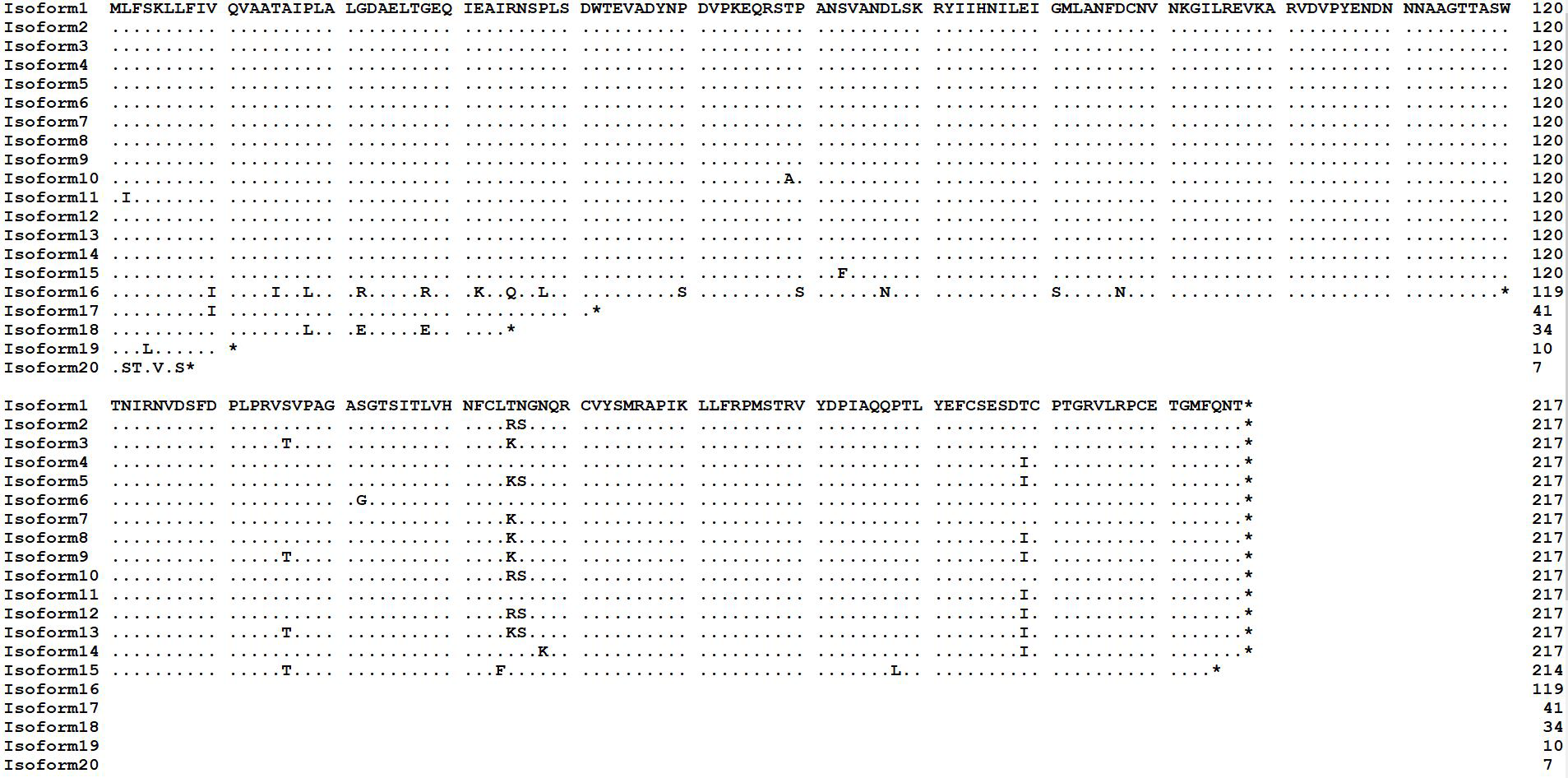
