## Supplemental Tables 1 and 2 for "The *Parastagonospora nodorum* necrotrophic effector SnTox5 targets the wheat gene *Snn5* and facilitates entry into the leaf mesophyll"

Supplementary Table 1. Primers used in this study

| ID | Sequence |
| --- | --- |
| Tox5HygDonor F1^*^ | **ATAGTCCATTGTCAGACTGGACAGAGGTCGCCGATTACAA**GTAAAACGACGGCCAG |
| Tox5HygDonor R1 | **AACGGAATTGGCAGGTGTAGAGCGCTGTTCTTTAGGGACG**CAGGAAACAGCTATGAC |
| Tox5_sgRNA | TTCTAATACGACTCACTATAGCTGTTCTTTAGGGACGTCTGTTTTAGAGCTAGA |
| SnTox5_pENTR_F1_bac | CACCATGGATGCGGAACTTACCGGAGAGCAAA |
| SnTox5_pENTR_R1 | CGTGTTCTGGAACATCCCTGTT |
| SnTox5_DONOR_F | GGGGACAAGTTTGTACAAAAAAGCAGGCTTTGTGCCTCTCATCAAATCG |
| SnTox5_DONOR_R | GGGGACCACTTTGTACAAGAAAGCTGGGTCCTACGTGTTCTGGAACATCC |
| Tox5_Seq_F | ACCCCAGTTTTGTCTTGATCC |
| M13F | GACGTTGTAAAACGACGGCCAGTG |
| M13R | CACAGGAAACAGCTATGACCATGA |
| HY | GGATGCCTCCGCTCGAAGTA |
| YG | CGTTGCAAGACCTGCCTGAA |
| PnActin_F | GCGGTGGCATCCACGTTACCACTTT |
| PnActin_R | TGCGATGATCTTGACCTTCATGGAC |
| SnTox5_qPCR_F | TGTAATGTCAACAAAGGCATCCTAC |
| SnTox5_qPCR_R | AGGCAAAAGTTGTGGACTAAAGTAA |

^*^Bolded sequence indicates the 40 bp region homologous to the flanking region of protospacer adjacent motif that used to disrupt *SnTox5*.

Supplementary Table 2. Phenotypic data used in GWAS analysis

| Isolate | Replicate 1^*^ | Replicate 2^*^ | Replicate 3^*^ | Average^*^ disease reaction |
| --- | --- | --- | --- | --- |
| 98-13025-2 | 1 | 0 | 0.5 | 0.5 |
| 98-13042-1 | 2 | 4 | 2.5 | 2.833333 |
| 98-13050-1 | 0 | 0.5 | 1.5 | 0.666667 |
| 98-13063-1 | 4.5 | 2.5 | 3.5 | 3.5 |
| 98-13066 | 3 | 4 | 2.5 | 3.166667 |
| 98-13082 | 4 | 4 | 3 | 3.666667 |
| 98-13091-2 | 3.5 | 3 | 2 | 2.833333 |
| AR1-1 | 3 | 4.5 | 3.5 | 3.666667 |
| AR2-1 | 3.5 | 3 | 2.5 | 3 |
| AR3-1 | 4 | 4.5 | 3 | 3.833333 |
| AR4-1 | 4 | 4.5 | 2 | 3.5 |
| AR5-1 | 0 | 0 | 0.5 | 0.166667 |
| AR6-1 | 2.5 | 2.5 | 1.5 | 2.166667 |
| BBC03 Sn-1 | 0 | 0 | 0.5 | 0.166667 |
| BBC03 Sn-3 | 0.5 | 1.5 | 0 | 0.666667 |
| BBC03 Sn-5 | 4 | 3.5 | 3.5 | 3.666667 |
| BBC04 Sn-1 | 1 | 2 | 3.5 | 2.166667 |
| BBC04 Sn-2 | 0.5 | 1.5 | 0 | 0.666667 |
| BBC04 Sn-5 | . | . | . | . |
| BBC04 Sn-6 | 0 | 1.5 | 2 | 1.166667 |
| FgoG10 Sn-1 | 0.5 | 0.5 | 2 | 1 |
| FgoN10 Sn-2 | 1.5 | 3.5 | 2 | 2.333333 |
| FgoN10 Sn-3 | . | 4 | 2 | 3 |
| FgoN10 Sn-4 | 2.5 | 2 | 1.5 | 2 |
| GA9-1 | 1 | 4 | 2.5 | 2.5 |
| GA9-2 | 4 | 4 | 4 | 4 |
| GA9-3 | 3 | 3 | 2 | 2.666667 |
| GA9-4 | 3.5 | 3 | 2.5 | 3 |
| GA9-5 | 1.5 | 1 | 0.5 | 1 |
| LDN03 Sn-1 | 2.5 | 3.5 | 4 | 3.333333 |
| LDN03 Sn-10 | 1.5 | 3.5 | 2.5 | 2.5 |
| LDN03 Sn-11 | 1 | 2.5 | 1.5 | 1.666667 |
| LDN03 Sn-2 | 4.5 | 2 | 4.5 | 3.666667 |
| LDN03 Sn-3 | . | 3.5 | 4 | 3.75 |
| LDN03 Sn-4 | . | 3.5 | 2 | 2.75 |
| LDN03 Sn-5 | 0.5 | 1.5 | 0.5 | 0.833333 |
| LDN03 Sn-6 | 1.5 | 1.5 | 2 | 1.666667 |
| LDN03 Sn-7 | 4 | 4.5 | 4.5 | 4.333333 |
| LDN03 Sn-8 | 2 | 2 | 2 | 2 |
| LDN03 Sn-9 | 3.5 | . | 1 | 2.25 |

*The “.” represents missing data.

Supplementary Table 2. Phenotypic data used in GWAS analysis

| Isolate | Replicate 1^*^ | Replicate 2^*^ | Replicate 3^*^ | Average disease reaction^*^ |
| --- | --- | --- | --- | --- |
| LDN05 SN-2 | 1.5 | 2.5 | 2 | 2 |
| LDN05 Sn-4 | 3 | 0.5 | 0 | 1.166667 |
| LDN05 Sn-5 | 0.5 | . | 0.5 | 0.5 |
| LDN07 Sn-1 | 0 | 2 | 0.5 | 0.833333 |
| LDN07 Sn-2 | 0 | 2.5 | 3 | 1.833333 |
| LDN07 Sn-3 | 0 | 0 | 0 | 0 |
| LDN07 Sn-4 | 0 | 1 | 0 | 0.333333 |
| LDN07Sn-5 | 0 | 1.5 | 2 | 1.166667 |
| LDN08Sn-1 | 0 | 0 | 0.5 | 0.166667 |
| LDN08Sn-2 | 0 | 1.5 | 0.5 | 0.666667 |
| LDN08Sn-3 | 2 | 1.5 | 2 | 1.833333 |
| LDN08Sn-4 | 2 | 0 | 2 | 1.333333 |
| LDN08Sn-5 | 0 | 3 | 1.5 | 1.5 |
| MD4-1 | 0.5 | 0.5 | 0.5 | 0.5 |
| MD4-2 | . | 2.5 | 0 | 1.25 |
| MD4-3 | 2 | 0 | 2 | 1.333333 |
| MN-2 | 0 | 0 | 0.5 | 0.166667 |
| MN-3 | 2.5 | 1.5 | 1.5 | 1.833333 |
| MN-4 | 0 | 1.5 | 0.5 | 0.666667 |
| MN-5 | 0 | 0 | 1.5 | 0.5 |
| MN-6 | 1.5 | 0 | 0 | 0.5 |
| MN-7 | 1.5 | 0 | 1.5 | 1 |
| MN-8 | 0 | 0 | 2 | 0.666667 |
| NC 7-1 | 0.5 | 1.5 | 0.5 | 0.833333 |
| NC 8-11 | 2.5 | 2.5 | 2.5 | 2.5 |
| NC 8-7 | 2 | 4 | 2.5 | 2.833333 |
| NC 9-12 | 2 | 3 | 3.5 | 2.833333 |
| NC 9-5 | 3 | 4.5 | 4 | 3.833333 |
| NC8-3 | . | 3 | 3 | 3 |
| NC8-4 | 2 | 3 | 3.5 | 2.833333 |
| NC8-6 | 2.5 | 2 | 2 | 2.166667 |
| NC8-8 | 4 | 3 | 4 | 3.666667 |
| NDJ16Sn-1 | 2 | 1.5 | 1.5 | 1.666667 |
| NDJ16Sn-10 | 3.5 | 2.5 | 1.5 | 2.5 |
| NDJ16Sn-11 | 3.5 | 1 | 1.5 | 2 |
| NDJ16Sn-12 | 2 | 2 | 3.5 | 2.5 |
| NDJ16Sn-13 | 2.5 | 0 | 4 | 2.166667 |
| NDJ16Sn-14 | 1.5 | 3 | 1.5 | 2 |
| NDJ16Sn-15 | 0 | 0 | 0 | 0 |
| NDJ16Sn-16 | 0 | 1.5 | 0 | 0.5 |

*The “.” represents missing data.

Supplementary Table 2. Phenotypic data used in GWAS analysis

| Isolate | Replicate 1^*^ | Replicate 2^*^ | Replicate 3^*^ | Average disease reaction^*^ |
| --- | --- | --- | --- | --- |
| NDJ16Sn-18 | 2.5 | 2.5 | 3 | 2.666667 |
| NDJ16Sn-19 | 2 | 0.5 | 3 | 1.833333 |
| NDJ16Sn-2 | 0.5 | 3 | 3 | 2.166667 |
| NDJ16Sn-20 | 0 | 0 | 2 | 0.666667 |
| NDJ16Sn-21 | 2 | 0 | 1.5 | 1.166667 |
| NDJ16Sn-22 | 4 | 0.5 | 2.5 | 2.333333 |
| NDJ16Sn-23 | 4 | 2 | 0.5 | 2.166667 |
| NDJ16Sn-24 | 2 | 1 | 2 | 1.666667 |
| NDJ16Sn-25 | 2.5 | 1 | 0.5 | 1.333333 |
| NDJ16Sn-3 | 1.5 | 0.5 | 2.5 | 1.5 |
| NDJ16Sn-4 | 4 | 2.5 | 3 | 3.166667 |
| NDJ16Sn-5 | 0.5 | 0 | 0 | 0.166667 |
| NDJ16Sn-6 | 0.5 | 2.5 | 3.5 | 2.166667 |
| NDJ16Sn-7 | 1.5 | 1 | 1 | 1.166667 |
| NDJ16Sn-8 | 3.5 | 3 | 2.5 | 3 |
| NDJ16Sn-9 | 1.5 | 1.5 | 1.5 | 1.5 |
| NDM16 Sn-1 | 1 | 2.5 | . | 1.75 |
| NDM16 Sn-10 | 3.5 | 2.5 | 2.5 | 2.833333 |
| NDM16 Sn-2 | 3 | 3.5 | 3.5 | 3.333333 |
| NDM16 Sn-3 | 1.5 | 3.5 | 0 | 1.666667 |
| NDM16 Sn-4 | 2.5 | 1.5 | 3 | 2.333333 |
| NDM16 Sn-5 | 4 | 3.5 | 4 | 3.833333 |
| NDM16 Sn-6 | 1 | 3 | 0.5 | 1.5 |
| NDM16 Sn-7 | 1.5 | 3 | 2.5 | 2.333333 |
| NDM16 Sn-8 | 1 | 0.5 | 1 | 0.833333 |
| NDM16 Sn-9 | 1 | 1.5 | 1.5 | 1.333333 |
| NDP16 SN-1 | 1.5 | 1 | 1.5 | 1.333333 |
| NDP16 SN-10 | 3 | 1.5 | 3 | 2.5 |
| NDP16 Sn-11 | 3 | 2.5 | . | 2.75 |
| NDP16 Sn-12 | 4 | 3.5 | 3 | 3.5 |
| NDP16 SN-2 | 0.5 | 0 | 0 | 0.166667 |
| NDP16 SN-3 | 0 | 0 | 0 | 0 |
| NDP16 SN-4 | 3.5 | 3.5 | 4 | 3.666667 |
| NDP16 SN-5 | 3 | 1.5 | 0.5 | 1.666667 |
| NDP16 SN-6 | 0.5 | 2 | 1.5 | 1.333333 |
| NDP16 SN-7 | 3.5 | 3 | 2.5 | 3 |
| OH03 Sn-1051 | 3 | 4 | 4 | 3.666667 |
| OH03 Sn-1180 | 3 | 2 | 4.5 | 3.166667 |
| OH03 Sn-123 | 3.5 | 3.5 | 3.5 | 3.5 |
| OH03 Sn-1354 | 2.5 | 3.5 | 4.5 | 3.5 |

*The “.” represents missing data.

Supplementary Table 2. Phenotypic data used in GWAS analysis

| Isolate^*^ | Replicate 1^*^ | Replicate 2^*^ | Replicate 3^*^ | Average disease reaction^*^ |
| --- | --- | --- | --- | --- |
| OH03 Sn-14 | 4 | 2.5 | 3.5 | 3.333333 |
| OH03 Sn-1501 | _. | 3 | 3.5 | 3.25 |
| OH03 Sn-1553 | 2.5 | 1.5 | 2.5 | 2.166667 |
| OH03 Sn-39 | 3 | 4.5 | 3.5 | 3.666667 |
| OH03 Sn-601 | 2.5 | 2.5 | 4.5 | 3.166667 |
| OH03 Sn-61 | _ | 0.5 | 2 | 1.25 |
| OH03 Sn-65 | 3 | 2.5 | 4.5 | 3.333333 |
| OH03 Sn-69 | 4 | 4 | 3 | 3.666667 |
| OH03 Sn-8 | 2.5 | 4 | 3.5 | 3.333333 |
| OH03 Sn-801 | 0.5 | 0.5 | 0 | 0.333333 |
| OH03 Sn-84 | 3.5 | 3 | 4.5 | 3.666667 |
| OH03 Sn-850 | 4.5 | 2 | 2 | 2.833333 |
| OKG16-1 | 3 | 2.5 | 3 | 2.833333 |
| OKG16-10 | 3 | 3 | 2 | 2.666667 |
| OKG16-11 | 3 | 3.5 | 3 | 3.166667 |
| OKG16-12 | 3.5 | 3.5 | 3.5 | 3.5 |
| OKG16-13 | 2 | 2 | 1.5 | 1.833333 |
| OKG16-14 | 2 | 1.5 | 1.5 | 1.666667 |
| OKG16-15 | 1.5 | 3.5 | 2 | 2.333333 |
| OKG16-16 | 2 | 1.5 | 3 | 2.166667 |
| OKG16-17 | 3.5 | 3.5 | 1 | 2.666667 |
| OKG16-2 | 4 | 3 | 2 | 3 |
| OKG16-3 | 4 | 2 | 2.5 | 2.833333 |
| OKG16-4 | 2 | 2.5 | 3 | 2.5 |
| OKG16-5 | 0.5 | 1.5 | 2.5 | 1.5 |
| OKG16-6 | 1.5 | 0 | 2 | 1.166667 |
| OKG16-7 | 4 | 3 | 2.5 | 3.166667 |
| OKG16-8 | 5 | 4.5 | 3.5 | 4.333333 |
| OKG16-9 | 2 | 2 | 2 | 2 |
| SC 3-1 | 2.5 | 3.5 | 3.5 | 3.166667 |
| SC 3-2 | 3.5 | 4 | 2.5 | 3.333333 |
| SC3-3 | 2 | 3 | 1.5 | 2.166667 |
| SC3-4 | 4 | 2 | 3.5 | 3.166667 |
| SDRF16-1 | 0 | 1 | 0 | 0.333333 |
| SDRF16-2 | 2.5 | 3 | 3 | 2.833333 |
| SDRF16-4 | 2 | 2 | 3 | 2.333333 |
| SDRF16-5 | 1.5 | 1.5 | 3.5 | 2.166667 |
| SDRF16-6 | 2 | 2 | 2.5 | 2.166667 |
| SDRF16-7 | 0 | 0.5 | 1 | 0.5 |
| SDRF16-8 | 0 | 0 | . | 0 |

*The “.” represents missing data.

Supplementary Table 2. Phenotypic data used in GWAS analysis

| Isolate^*^ | Replicate 1^*^ | Replicate 2^*^ | Replicate 3^*^ | Average disease reaction^*^ |
| --- | --- | --- | --- | --- |
| SDRF16-9 | 0.5 | 1 | 0 | 0.5 |
| Sn2000 | 4.5 | 3 | 4.5 | 4 |
| SN330NY91 | 1.5 | 2 | 2 | 1.833333 |
| SN335NY91 | 2 | 1.5 | 0 | 1.166667 |
| SN345NY91 | 1 | 0 | 0 | 0.333333 |
| SN349NY91 | _ | 4 | 1.5 | 2.75 |
| SN351NY91 | 2.5 | 2.5 | 3 | 2.666667 |
| SN356NY91 | 0 | 1 | 1 | 0.666667 |
| SN358NY91 | 3 | 1.5 | 2.5 | 2.333333 |
| SN366NY91 | 0 | 3 | 1.5 | 1.5 |
| SN369NY91 | 0 | 0 | 0 | 0 |
| SN377NY91 | 4 | 0.5 | 2 | 2.166667 |
| Sn50 | 2 | 3 | 4.5 | 3.166667 |
| Sn-6 (Sn69-1) | 0.5 | 0 | 3.5 | 1.333333 |
| SnOre11-1 | 3.5 | 2.5 | 3 | 3 |
| SnOre11-2 | 3.5 | 3 | 3.5 | 3.333333 |
| SnOre11-3 | 2.5 | 3.5 | 3.5 | 3.166667 |
| SnOre11-4 | . | 4 | 3 | 3.5 |
| SnOre11-5 | 3.5 | 3.5 | 3 | 3.333333 |
| SnOre11-6 | 3 | 3.5 | 4.5 | 3.666667 |
| SnOre11-7 | 4.5 | 2.5 | 3.5 | 3.5 |
| SnOre11-8 | 4 | 2 | 2.5 | 2.833333 |
| SNOV92X D1.3 | 2.5 | 2.5 | 3 | 2.666667 |
| SNOV92X D4.1 | 3.5 | 3.5 | 2 | 3 |
| SNOV92X F1.1 | 4 | 2 | 2 | 2.666667 |
| SNOV92X F2.1 | 3 | 3 | 2.5 | 2.833333 |
| SNOV92X H1.4 | 3 | 3 | 3.5 | 3.166667 |
| TN 5-1 | 4.5 | 5 | 3.5 | 4.333333 |
| TN5-2 | 1.5 | 0 | 0.5 | 0.666667 |
| TN5-3 | . | 0 | 1 | 0.5 |
| TN5-4 | 3.5 | 2.5 | 2.5 | 2.833333 |
| TN5-5 | 4 | 3 | 3 | 3.333333 |
| VA 5-2 | 2 | 2.5 | 3 | 2.5 |
| VA 5-3 | 3 | 3.5 | 4 | 3.5 |
| VA 5-4 | 3.5 | 3 | 3 | 3.166667 |
| VA 5-5 | 4 | 0.5 | 3.5 | 2.666667 |

*The “.” represents missing data.
